# Primate superior colliculus is engaged in abstract higher-order cognition

**DOI:** 10.1101/2023.01.17.524416

**Authors:** Barbara Peysakhovich, Stephanie M. Tetrick, Alessandra A. Silva, Sihai Li, Ou Zhu, Guilhem Ibos, W. Jeffrey Johnston, David J. Freedman

**Affiliations:** Department of Neurobiology, University of Chicago, Chicago, IL, 60637; Institut de Neurosciences de la Timone, Aix-Marseille Université, CNRS, Marseille, France, 13005; Neuroscience Institute, University of Chicago, Chicago, IL, 60637

**Keywords:** superior colliculus, cognition, categorization, subcortical, vision, parietal, decision making, primate, electrophysiology, reversible inactivation

## Abstract

Categorization is a fundamental cognitive process by which the brain assigns stimuli to behaviorally meaningful groups. Investigations of visual categorization in primates have identified a hierarchy of cortical areas that are involved in the transformation of sensory information into abstract category representations. However, categorization behaviors are ubiquitous across diverse animal species, even those without a neocortex, motivating the possibility that subcortical regions may contribute to abstract cognition in primates. One candidate structure is the superior colliculus (SC), an evolutionarily conserved midbrain region that, although traditionally thought to mediate only reflexive spatial orienting, is involved in cognitive tasks that require spatial orienting. Here, we reveal a novel role of the primate SC in abstract, higher-order visual cognition. We compared neural activity in the SC and the posterior parietal cortex (PPC), a region previously shown to causally contribute to category decisions, while monkeys performed a visual categorization task in which they report their decisions with a hand movement. The SC exhibits stronger and shorter-latency category encoding than the PPC, and inactivation of the SC markedly impairs monkeys’ category decisions. These results extend SC’s established role in spatial orienting to abstract, non-spatial cognition.

## Introduction

The superior colliculus (SC), a brainstem region that is evolutionarily conserved across vertebrates, has long been known to play a crucial role in directing orienting movements of the eyes and head [1–11]. Although traditionally thought to be a reflexive structure that receives and implements motor instructions from upstream brain areas, the SC is involved in the selection of which target to orient to [12–15], and has been shown to be engaged in complex behavioral tasks that involve either overt target selection, such as decision-making tasks in which animals report their choice with a saccade to a particular target [16–23], or covert target selection, such as tasks that require animals to attend to stimuli at particular spatial locations [24–31]. However, it is unknown whether the involvement of the SC in cognitively demanding tasks is restricted to contexts involving spatial orienting or target selection.

Here, we investigated whether the primate SC is more generally involved in abstract cognition. We trained monkeys to perform an abstract visual categorization task that dissociates sensory, cognitive, and motor components, and compared neuronal activity in the SC and the lateral intraparietal area (LIP), a cortical region in the posterior parietal cortex that is anatomically interconnected with the SC [32–37] and has been previously shown to causally contribute to category processing [38]. We also assessed the causal contribution of the SC to category decisions using reversible pharmacological inactivation. We show that the SC exhibits robust, short-latency encoding of abstract categories and that inactivation of the SC markedly impairs animals’ categorization task performance. These results indicate that the primate SC plays an unexpected key role in higher-order cognition, independent of its role in spatial orienting. In addition, we show that category and saccade-related signals are encoded in near-orthogonal subspaces in population activity in the SC, providing an explanation for how a motor structure like the SC can be recruited to participate in more flexible cognitive behaviors.

## Results

### Behavior

Two monkeys performed a delayed match-to-category (DMC) task in which they grouped 360° of motion directions into two categories based on an arbitrary category rule. The categories were defined by two perpendicular boundaries that produced four 90°-wide quadrants (**Fig. 1a**). To disambiguate neuronal encoding of direction vs. category, opposite quadrants were assigned to the same category, so that motion directions that are 180° apart belonged to the same category while nearby directions were often in different categories. Monkeys viewed sample and test dot-motion stimuli separated by a 1.2 sec memory delay (**Fig. 1b**). On each trial, they received a fluid reward for releasing a manual touch bar when the category of the test stimulus matched the category of the sample stimulus (Match trials). If the test stimulus category did not match the sample stimulus category (Non-match trials), the monkeys were shown a second test stimulus, which always matched the sample category (and required release of the touch bar in order to receive a reward). The monkeys were required to maintain gaze on a central fixation spot throughout the trial (see *Methods*). The monkeys’ decisions about the sample category were abstract because the two categories were defined by the learned arbitrary boundaries, and because they were not linked to different motor actions or plans.

**Fig. 1:**
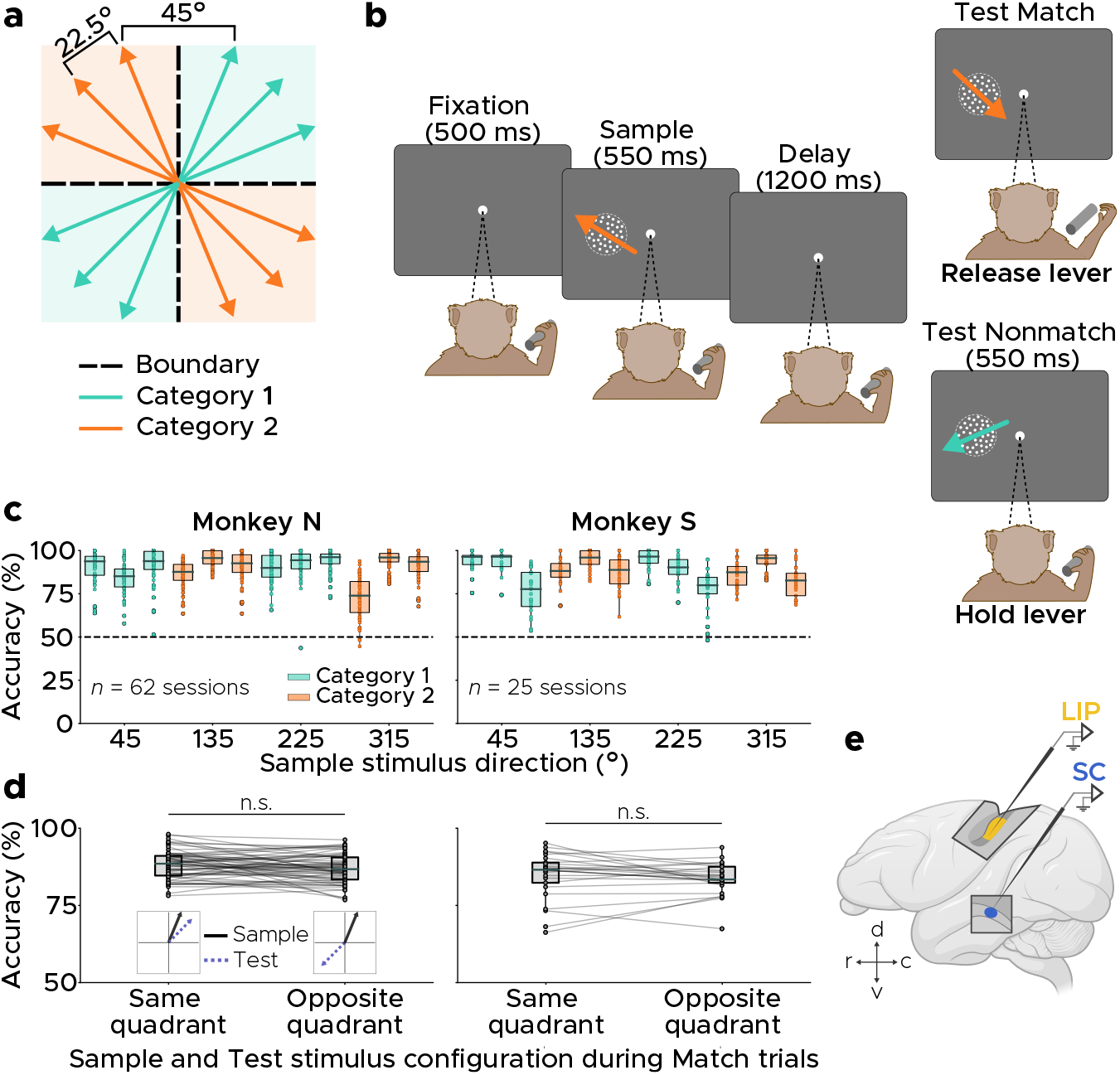
Monkeys learn to categorize motion stimuli based on an arbitrary category rule. **a,** Stimulus geometry of the two-boundary delayed match-to-category (DMC) task. 12 directions of motion are grouped into two categories based on two orthogonal category boundaries (dashed lines), such that directions that are 180° apart belong to the same category. Directions within the same quadrant are 22.5° apart, and near-boundary directions are 22.5° from the boundary. **b,** Trial structure of the DMC task. Monkeys were required to maintain gaze fixation within a small window centered on a central fixation cue, and reported their decisions with a hand movement. **c,** Behavioral performance across recording sessions for each of the 12 sample stimulus directions for Monkey N (left) and Monkey S (right). Horizontal dashed line indicates chance performance. **d,** Behavioral performance across sessions on Match trials in which the sample and test stimuli were in the same or opposite quadrants. There was no significant difference in mean performance on trials in which sample and test stimuli were in the same or different quadrants (Monkey N: Same quad. = 88.2 ± 4.6%, Opp. quad. = 86.9 ± 4.70%, *P* = .094, Monkey S: Same quad. = 84.9 ± 7.4%, Opp. quad. = 83.8 ± 5.2%, *P* = .580, permutation test). **e**, Schematic of neural recording locations in the lateral intraparietal area (LIP) and superior colliculus (SC).

Following long-term training, both monkeys performed the DMC task with >85% mean accuracy during neural recording sessions (Monkey N: 89.40 ± 3.43%, Monkey S: 88.20 ± 3.34%; **Fig. 1c**), with no significant difference in mean performance between LIP and SC recording sessions (Monkey N: *P* = .094, Monkey S: *P* = .229, permutation test; **Extended Fig. 1)**. Monkeys performed similarly on Match trials in which sample and test stimuli were in the same vs. opposite quadrants (Monkey N: *P* = .094, Monkey S: *P* = .580, permutation test; **Fig. 1d**), indicating that they learned to categorize stimuli across opposite quadrants.

### Robust encoding of sample category in the SC

We analyzed spiking activity during the DMC task from 555 LIP neurons (Monkey N: 228, Monkey S: 327) and 604 SC neurons (Monkey N: 362, Monkey S: 242; **Fig. 1e**). Individual LIP neurons often showed binary-like category selectivity during the sample and delay task periods, with distinct activity for directions in different categories and similar activity for directions in the same category (**Fig. 2a**), consistent with previous studies that used a similar categorization task with a simpler linear category boundary [39–43]. This category selectivity extended even to stimuli in opposite quadrants that belong to the same category. Remarkably, individual SC neurons also showed strong category selectivity during sample stimulus presentation and the subsequent delay and test periods (**Fig. 2b**).

**Fig. 2:**
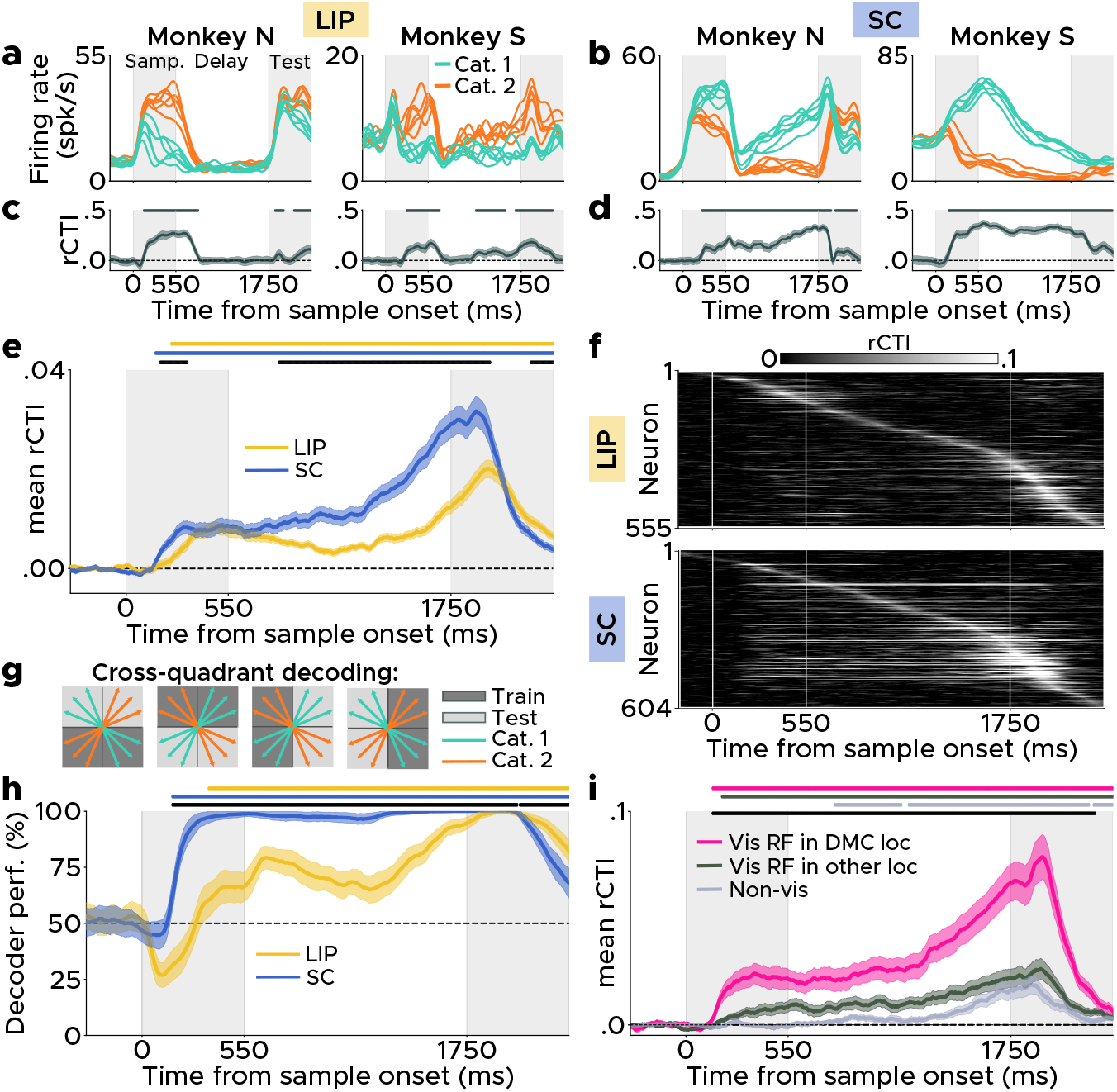
Neural activity in the SC contains reliable and short-latency sample stimulus category information. **a,** Peristimulus time histograms of example category-tuned LIP neurons. **b,** Same as **a**, but for SC. **c,** ROC-based category tuning index (rCTI) across time in trial for the two example LIP neurons in **a**. Shading indicates s.d., computed via resampling of trials. Grey symbols at the top of the plots indicate time points at which rCTI is significantly above chance. **d,** Same as **c**, but for SC. **e,** Time course of mean rCTI across LIP and SC neurons. Shading indicates s.e.m. **f,** Matrix of rCTI for all neurons in LIP and SC, where each row shows a single neuron’s rCTI as a function of time in the trial. **g,** Schematic of cross-quadrant sample category classifiers, which are trained on trials from two quadrants (dark gray) and validated on trials from the remaining two quadrants (light gray). **h,** Time course of mean accuracy of cross-quadrant category classifiers for LIP and SC populations. Shading indicates s.d. **i,** Time course of mean rCTI across SC neurons that are visually responsive to stimuli at locations that overlap with the position of the DMC stimuli (pink), neurons that are visually unresponsive at DMC locations but visually responsive at other locations (dark grey), and neurons that are visually unresponsive (light grey). In **e**, **h** and **i**, colored symbols above plots indicate timepoints at which values significantly exceed chance. In **e** and **h**, black symbols above panel indicate time points at which there is a significant difference between areas, and in **i**, black symbols above panel indicate time points at which mean rCTI of Vis neurons is significantly higher than both Vis-other and Non-vis neurons (*P* < .050, two-tailed permutation tests).

We quantified the strength, trial-by-trial reliability, and time course of category tuning in individual neurons using an ROC-based category tuning index (rCTI), which compares neuronal discrimination between pairs of directions in the same vs. different categories (**Extended Fig. 2a**; see *Methods*). rCTI values can range from −0.5 to 0.5, with positive values indicating larger differences in firing rates between pairs of directions in different vs. same categories (and thus strong category tuning) and negative values indicating the opposite. **Fig. 2c** and **d** show the time course of rCTI for the single-neuron examples in **Fig. 2a** and **b**, and **Fig. 2f** shows a heatmap of rCTI values for all neurons in LIP and SC. We classified neurons as category-tuned if they showed significantly elevated rCTI values relative to a null rCTI distribution for at least 30 consecutive ms; **Extended Fig. 2;** see *Methods*).

In the LIP, 69.5% of neurons were category-tuned (Monkey N: 80.7%, Monkey S: 61.8%), and in the SC, 60.3% of neurons were category-tuned (Monkey N: 66.0%, Monkey S: 51.7%). In both LIP and SC, mean rCTI across neurons was shifted toward positive values at nearly every time point of the DMC task following sample onset (**Fig. 2e**). The increase in mean rCTI above baseline levels during the sample period of the task occurred significantly earlier in the SC than in the LIP (LIP: 245 ms, SC: 160 ms, *P* = .008, permutation test; **Fig. 2e**, yellow and blue symbols above panel). Moreover, rCTI values were significantly greater in the SC than the LIP throughout much of the trial (**Fig. 2e**, black symbols above panel), indicating stronger category encoding in the SC than LIP. During the sample epoch, a significantly greater percentage of LIP neurons were direction tuned compared to SC (LIP: 30.2%, SC: 20.4%; *X*^2^ = 7.65, *P* = .006, *X*^2^ test). To determine whether the earlier and stronger category tuning in the SC compared to LIP can be explained by the difference between areas in the proportion of direction-tuned neurons, we computed the mean rCTI in each area only on direction-untuned neurons (see *Methods*). This analysis again revealed shorter-latency category selectivity in SC compared to LIP (LIP: 195 ms, SC: 155 ms; *P* = .042, permutation test).

We next used linear support vector machine (SVM) classifiers to quantify the amount and timing of direction-independent category encoding in the LIP and SC neural populations. To evaluate the strength of category encoding in a direction-independent manner, the classifiers were trained on trials from two quadrants (one from each category) and validated on the remaining two quadrants (**Fig. 2g**). The logic behind this classifier is that, if the neural populations robustly encode category in a binary-like format, the classifier will be able to generalize between the two quadrants within the same category. In addition, this approach prevents direction tuning from contributing to category decoding performance by decorrelating direction and category between the sample and test sets; note that we find below-chance classifier performance when the population shows strong direction tuning. We also assessed the strength and time course of motion direction encoding in the neural populations from both brain areas using a category-independent direction classifier, as shown in **Extended Fig. 3**.

In the SC, category classifier accuracy rapidly increased above chance levels within approximately 170 ms of sample stimulus onset, and remained at almost 100% throughout the rest of the trial (**Fig. 2h**). In the LIP, category classifier accuracy was below chance shortly after sample stimulus onset, indicating that direction tuning was more dominant than category tuning during the early sample epoch. Sample category information could be decoded more reliably from SC than LIP activity throughout the sample, delay, and early test phases of the task (**Fig. 2h**, black symbols above panel), indicating stronger sample category encoding in the SC neural population than the LIP population.

Results regarding the strength and timing of category selectivity for the rCTI and decoding analyses were qualitatively similar in the two animals (**Extended Fig. 4**). The remarkably strong and short-latency encoding of abstract category in the SC is surprising, since primate SC is strongly associated with oculomotor control, orienting, and target selection, as opposed to higher-order non-spatial cognitive functions like categorization, and the animals here had to maintain central gaze fixation and did not report their decisions with saccadic eye movements.

### Category encoding in the SC cannot be explained by eye movements

Given SC’s well-established role in directing saccadic eye movements [1–11], one explanation for the unexpected presence of category selectivity in the SC is that the reported category signals could be a result of distinct patterns of microsaccades during different conditions of the DMC task. Interestingly, we observed that the monkeys produced idiosyncratic, category-specific eye movements (within the allowed fixation window) that were highly stereotyped across sessions (**Extended Fig. 5**). The category-specific microsaccades occurred only during the memory delay epoch in Monkey N and primarily during the memory delay epoch in Monkey S, indicating that the monkeys’ eye movements reflect their working memory contents, consistent with results from past work in monkeys performing a delayed matching task [44]. To understand whether these eye movements can explain the observed category signals in the SC, we first compared the time course of neuronal category selectivity in the SC and the time course of category-specific eye position. This revealed that the two time courses were highly decoupled in time (**Extended Fig. 6a**), with category selectivity preceding category-specific eye position by several hundred milliseconds (Monkey N: neural decoder = 175 ms, eye decoder = 1075 ms; Monkey S: neural decoder = 170 ms, eye decoder = 865 ms). We next built linear encoding models [45] to determine whether firing rates of individual neurons (across trials and time within trial) are better predicted by stimulus category or eye movements (see *Methods*). In the majority of SC neurons, firing rates during the DMC task were better predicted by the stimulus than by eye movements (**Extended Fig. 6f**). These results indicate that category selectivity in the SC cannot be accounted for by the category-specific eye movements during the DMC task, and raise the possibility that the eye movements may rather be a consequence of the presence of category selectivity in the SC.

### Preferential encoding of category in visual SC neurons

The SC is a core stage of oculomotor processing and contains a diversity of neuronal response types based on firing rate modulation to visual, visuomotor, and motor aspects of visually- and memory-guided saccade (MGS) tasks. We were interested in understanding whether abstract category encoding in the SC was more prevalent among SC neurons with particular patterns of visual or motor selectivity. At the beginning of each DMC recording session, monkeys performed a block of the MGS task (**Extended Fig. 7a,b**), allowing a comparison of neuronal activity and selectivity from the same neurons during the two tasks. We analyzed activity from a subset of 423 SC neurons (Monkey N: 259, Money S: 164) from which we recorded during both the DMC and MGS tasks. Examples of single-neuron responses from SC during the MGS task are shown in **Extended Fig. 7c** and **d**).

The short-latency category encoding in the SC raises the possibility that the SC plays a direct role in the rapid bottom-up categorization of incoming visual stimuli (i.e., the transformation of direction information into category representations). One piece of evidence that would support such a role is if the category signal first emerges in visually-responsive neurons whose receptive fields match the position of the DMC stimuli. We therefore examined whether SC category selectivity varied between three groups of neurons: (1) those that are visually responsive during the MGS task to stimuli presented at locations that overlap with the stimulus position in the DMC task (Vis neurons), (2) neurons that are visually unresponsive at the DMC stimulus location but visually responsive at other locations (Vis-other), and (3) visually unresponsive neurons (Non-vis; see *Methods* for details). We compared mean rCTI for 115 Vis neurons (Monkey N: 82, Monkey S: 33), 178 Vis-other neurons (Monkey N: 97, Monkey S: 81), and 130 Non-Vis SC neurons (Monkey N: 80, Monkey S: 50). Category selectivity emerges significantly earlier in Vis neurons than in Vis-other neurons (Vis: 145 ms, Vis-other: 195 ms, *P* = .036, permutation test) or Non-vis neurons (Non-vis: 800 ms, *P* < .001, permutation test; **Fig. 2i**), and mean rCTI is significantly higher in Vis neurons compared to the other two groups throughout much of the trial (**Fig. 2i**, black symbols above panel).

### Orthogonal encoding of saccades and stimulus category in the SC

Next, we sought to understand the structure of population activity that allows motor and non-motor representations to coexist within a core oculomotor region like the SC. How is it that the SC can be strongly modulated by stimulus category (or visual information in general) without producing task-interfering saccades, given that injection of even a small amount of electrical current into intermediate and deep layers of the SC can reliably generate large-amplitude eye movements [5, 6]? One explanation is that independent populations of SC neurons might participate in saccade and category encoding. To investigate this possibility, we characterized the amount of overlap in neural populations that are category-selective during the DMC task and saccade direction-selective during the MGS task (see *Methods*). We observed substantial overlap in the population of SC neurons that are category-selective during the DMC task and saccade-direction-selective during the MGS task (**Fig. 3a**). In addition, we compared neuronal responses during the peri-saccade period of the MGS task and sample period of the DMC task, and calculated a task-preference index to quantify the ratio of how much each neuron is modulated by the two tasks (see *Methods*). If there are separate populations of neurons that encode saccade parameters and category, the distribution of task-preference indices across neurons would be bimodal. However, this was not observed in the data: the distributions peaked near zero, indicating that a majority of neurons participated in both tasks (**Fig. 3a**), with no evidence for bimodality (Hartigan’s dip test, Monkey N: *P* = 0.984, Monkey S: *P* = 0.988).

**Fig. 3:**
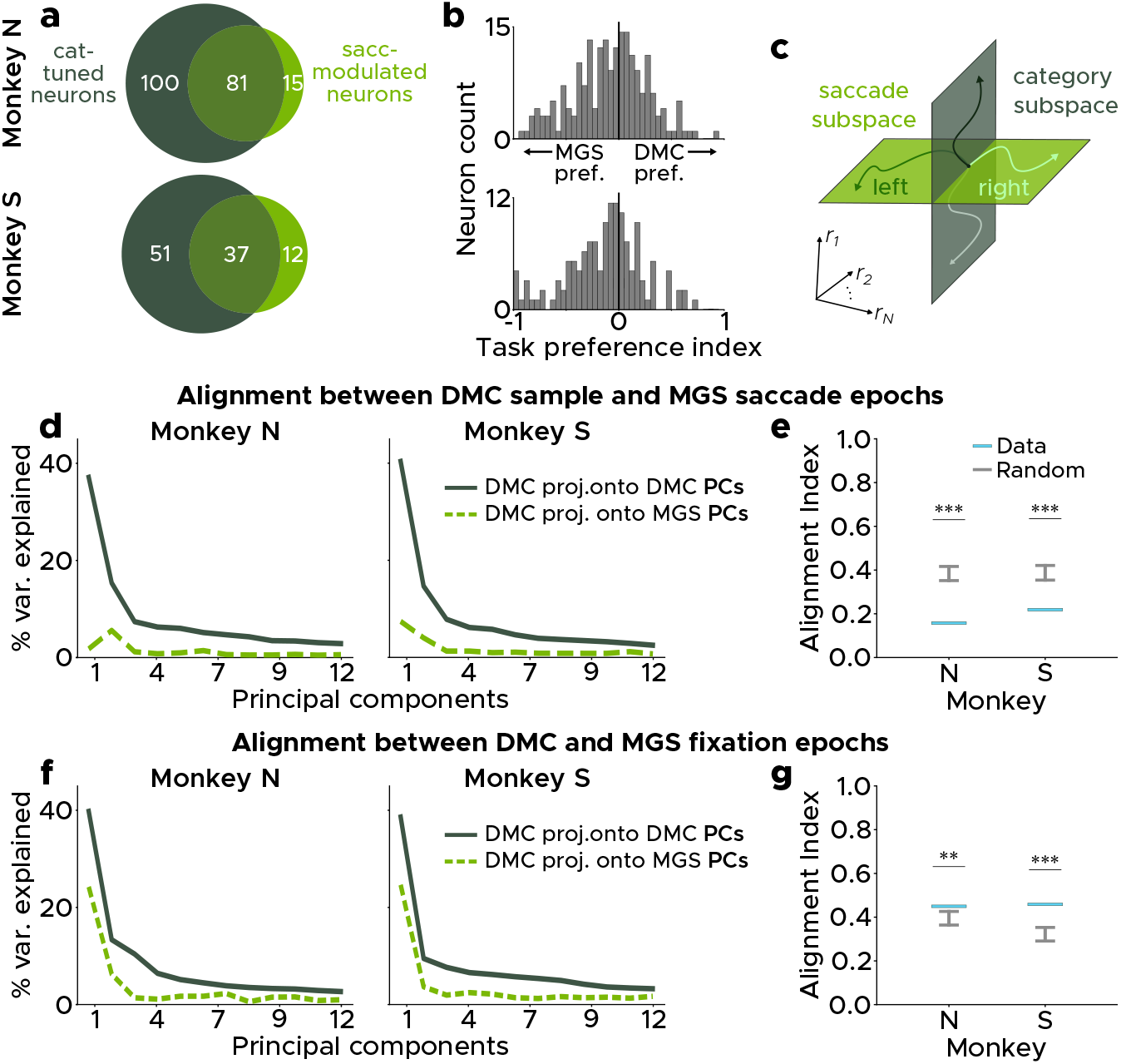
Orthogonal population-level encoding of saccade and category in the SC. **a,** Venn diagrams showing the overlap of neurons that are category-tuned during the DMC task (dark green) and neurons that are saccade-direction-modulated during the MGS task (light green; see *Methods*). **b,** Distributions of task preference indices across neurons for Monkey N (bottom) and Monkey S (top). Positive values indicate preference for the DMC task, and negative values indicate preference for the MGS task. **c,** Schematic of hypothesized orthogonal saccade and category activity subspaces. **d,** The percentage of variance of DMC sample period data explained when projected onto its own top 12 principal components (solid dark green line) or onto the top 12 principal components defined by activity during the peri-saccade period (−200 to +25 from saccade onset) of the MGS task (dotted light green line). **e**, Alignment index between the DMC sample epoch data and MGS peri-saccade data. The alignment index, which is the ratio of the two traces shown in **d**, equals 1 when two subspaces are perfectly aligned and equals 0 when two subspaces are perfectly orthogonal. Blue: alignment indices for the real data. Grey: 95% confidence intervals of alignment indices between pairs of random vector projections from data (see *Methods*). For both monkeys, the real data are significantly more orthogonal than expected by chance (*P* < .001). **f** and **g**, same as **d** and **e** but for the fixation epochs (−500 to 0 ms relative to stimulus onset) for the DMC and MGS tasks. The data are significantly more aligned than expected by chance (Monkey N: *P* = .004, Monkey S: *P* < .001).

Another possibility is that the structure of population activity in the SC is organized to maintain approximately orthogonal neural representations during category processing and saccade planning, such that projection of category-related neural activity onto the saccade-encoding neural axis produces minimal eye movements (**Fig. 3c**). To investigate this idea, we quantified the degree of alignment between the neural activity space from the sample period of the DMC task (from +150 to +550 ms relative to sample onset) and the peri-saccade period of the MGS task (from −200 to 0 ms relative to saccade onset). This approach [46] compares the percentage of DMC data variance explained when the DMC data are projected onto DMC-defined vs. MGS-defined neural axes, and produces a single alignment index (AI) value that ranges from 0 (indicating perfect orthogonality between the two subspaces) and 1 (indicating perfect alignment). To determine whether the resulting indices are more or less aligned than random, we compared the AI computed from the data to a null distribution of AI values between subspaces drawn from a random space that shares a covariance structure with the real data (see *Methods*).

In both monkeys, alignment between the category and saccade subspaces was near-orthogonal; projection of the DMC data onto the MGS axes captured minimal DMC data variance (**Fig. 3d**), and the resulting alignment index was significantly closer to 0 than expected by chance (**Fig. 3e**; Monkey N: AI = .159, *P* < .001, Money S: AI = .222, *P* < .001). This result is unlikely due to neural fluctuations over time in the session; in both monkeys, DMC activity from the beginning of the sessions was closely aligned to DMC activity from end of the sessions (**Extended Fig. 8a**), and alignment to MGS activity was similarly low for DMC activity from the beginning vs end of the sessions (**Extended Fig. 8b**).

To determine whether this misalignment between category and saccade subspaces may be due to general differences in behavioral state in the two task contexts, we quantified the degree of alignment between the baseline neural activity during the fixation epochs (from −500 to 0 ms relative to stimulus onset) for the two tasks. During this baseline period, the task demands (i.e., maintaining fixation) are shared between the two tasks, but the overall behavioral context is different. In both monkeys, fixation epoch activity during the DMC and MGS tasks was significantly more aligned than expected by chance (**Fig. 3f-g**; Monkey N: AI = .451, *P* = .007, Money S: AI = .460, *P* < .001), indicating that the misalignment between category and saccade subspaces cannot be explained by differences in behavioral state between the two tasks, and suggesting that the SC may selectively use an orthogonal-coding strategy to minimize motor interference.

Together, these results suggest a mechanism by which neural populations in the SC can multiplex motor signals and the higher-order, non-motor cognitive signals that we report here. The SC may use a similar mechanism to encode visual information during contexts in which animals do not need to produce (or are required to withhold) eye movements. These results also provide a possible explanation for the stereotyped, category-specific microsaccades that emerge several hundred ms after neural category selectivity onset during the DMC task (**Extended Fig. 5**); these category-specific eye movements may reflect “leak” from the category subspace to the saccade subspace. During learning of the DMC task, the SC network may arrive at a solution (i.e., a particular geometry of population activity) that is sufficiently (though not perfectly) orthogonal to the saccade subspace, such that any resulting eye movements are within the behavioral constraints of the task (i.e., fall within the allowed fixation window).

### Reversible pharmocological inactivation of the SC impairs performance on DMC task

Finally, we sought to determine whether category-correlated neural activity in the SC plays a causal role in the DMC task by infusing muscimol, a GABA_A_ agonist, to reversibly inactivate the SC (**Fig. 4a**). We compared monkeys’ behavior during SC inactivation with that during control sessions collected on the same day before injection.

**Fig. 4:**
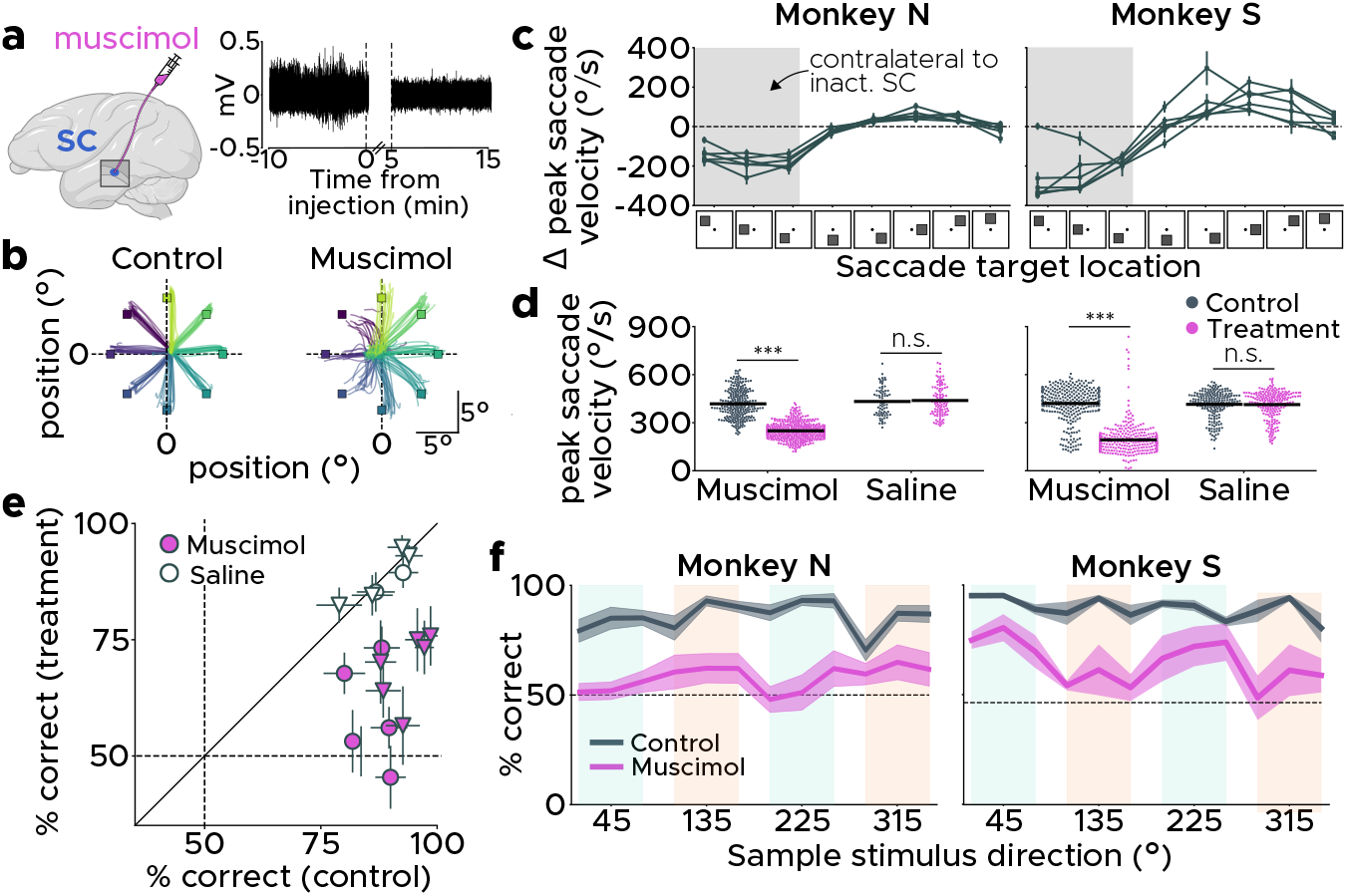
SC is causally involved in categorization task performance. **a,** We reversibly inactivated the SC using muscimol, a GABA_A_ agonist (left). Shortly following injection of muscimol, spiking activity markedly decreased in the surrounding tissue (right). **b,** Trajectories of eye movements in Monkey S during the visually-guided saccade task for an example session before (left) and after (right) muscimol injection. Individual traces represent eye gaze trajectories for individual trials and are color-coded by condition. Small colored squares indicate the position of saccade targets for different conditions. **c,** Changes in peak saccade velocity during the memory/visually-guided saccade task before and after SC inactivation at each of the eight stimulus locations used in the task. Grey background shading indicates conditions in which the target location was in the inactivated hemifield. **d,** Peak saccade velocity for trials (across all sessions) in which the target was in the inactivated hemifield during control blocks (black) and treatment blocks (pink) for muscimol and saline injection sessions. Horizontal black lines indicate mean of distributions. **e,** Overall session accuracy for the DMC task before vs. during SC inactivation. Unfilled: saline injection, Filled: muscimol, Circles: Monkey N, Triangles: Monkey S. Error bars: 95% multinomial confidence intervals for each session. **f,** Mean performance on the DMC task on each of the 12 sample motion stimuli during control behavior (dark grey) and SC inactivation (pink). *** *P* < .001, permutation test.

We first verified the efficacy of SC inactivation by monitoring changes in saccade velocity for saccades towards the inactivated hemifield during a memory-guided (Monkey N) or visually-guided (Monkey S) saccade task. With the small dose of muscimol injected into SC (see **Extended Table 1**), both monkeys were able to successfully perform the memory/visually-guided saccade task following injection, even for targets in the inactivated hemifield. However, both monkeys exhibited a large reduction in accuracy and peak velocity of saccades to targets contralateral to the inactivated hemisphere (**Fig. 4b-d**), an effect that is characteristic of reversible SC inactivation [9–11]. This difference in mean peak velocity between the control and post-treatment blocks was significant for data combined across all muscimol injection sessions (Monkey N: control = 418 ± 90°/s, treatment = 249 ± 51°/s, *P* < .001; Monkey S: control = 420 ± 106°/s, treatment = 193 ± 92°/s, *P* < .001, permutation test; **Fig. 4d**), and was consistent (and significant) for all individual muscimol injection sessions (**Extended Table 2**). This effect was absent for data across sessions in which the SC was infused with saline (Monkey N: control = 432 ± 90°/s, treatment = 438 ± 95°/s, *P* = .673, Monkey S: control = 415 ± 84°/s, treatment μ= 413 ± 87°/s, *P* = .848, permutation test) and for all individual saline injection sessions (**Extended Table 2**).

Both monkeys showed a marked impairment in DMC task performance during muscimol injection sessions (**Fig. 4e-f**). We observed a significant reduction in accuracy on the DMC task in every experimental session in which SC was injected with muscimol, and no change in accuracy for saline injection sessions (**Extended Table 3**). These results are consistent with SC being causally involved in performance of the DMC task.

The behavioral impairment on the DMC task during SC inactivation is unlikely to be entirely due to deficits in low-level visual processing of stimuli presented in the inactivated hemifield. The DMC deficit is not purely an attentional/hemispatial neglect-like impairment, as the monkeys are still able to perceive, attend to, and respond to stimuli presented in the inactivated hemifield during the MGS task, and still attempt the DMC task and respond at appropriate times in the trial (i.e., release the lever only during the test epochs). The DMC deficit is also unlikely due to an impairment in the sensory processing of motion stimuli, as previous studies using similar experimental protocols have shown that SC inactivation produces only very slight impairments in direction discrimination of high-coherence motion stimuli like those used in our DMC task [28].

It is important to note that our experimental design cannot isolate the precise nature of the deficit caused by SC inactivation, as the DMC task requires several complex computations, including the transformation of sample direction into category, the maintenance of sample category information in working memory, the computation of the test stimulus category, and the comparison of sample and test stimulus categories. Future experiments with additional control tasks can more precisely characterize the nature of the deficit(s) caused by SC inactivation during this task. Despite this limitation, our study reveals that the functions of primate SC extend beyond those ascribed to it by current models.

## Discussion

Our results demonstrate that the role of the primate SC extends beyond sensorimotor functions and spatial orienting to abstract, higher-order cognitive processing, even in a task that does not involve reporting decisions with saccades. The SC robustly encodes the learned categories of visual stimuli during all DMC task phases (including stimulus presentation, short-term memory, and comparison periods), and category signals in the SC arose with a shorter latency and were stronger than in LIP, an area previously shown to be causally involved in a similar categorization task. Moreover, reversible inactivation of the SC markedly impairs category task performance, indicating that activity in the SC is causally involved in abstract categorization.

Our findings also suggest that the SC neural population multiplexes category and saccade information by projecting those variables into distinct and near-orthogonal activity subspaces. This can explain how a motor structure like the SC can encode other task variables without producing task-interfering eye movements, closely related to the mechanism proposed to explain motor planning activity (without motor output) in the primary motor cortex [46]. Even more broadly, this could be a general principle of neural coding through which a single neural population can efficiently and robustly encode multiple factors.

Previous studies that observed cognitive and/or abstract encoding in the SC used tasks in which animals either spatially orient to a particular target to indicate their choices [16–23, 47–49], or tasks in which animals need to covertly orient to stimuli at distinct cued locations in different task conditions [24–31]. By contrast, in the current study, monkeys did not report their decisions with a saccade or orient attention to different locations for different categories. Thus, it is difficult to account for our results based on differences in covert spatial attention. The DMC task also required monkeys to maintain gaze fixation throughout the trial, and category encoding in SC could not be explained by the animals’ patterns of eye movements during the task.

We investigated SC during the DMC task because of evidence that cortical areas that are closely involved in oculomotor functions, such as LIP and the frontal eye fields (FEF), are also engaged in abstract categorization and flexible decision tasks [50]. Our previous work shows that LIP plays a causal role in abstract categorization [38], and that it preferentially encodes motion categories compared both to visual cortical areas MT and MST [39, 51], and executive control areas such as the lateral prefrontal cortex [40]. Anatomical connections between LIP and SC also motivated examining SC [32–37], as well as the similar patterns observed in SC and LIP during saccade-based tasks. We were also inspired by work showing that SC activity reflects higher-order functions such as attention and perceptual decisions [16–31, 47]. Our results highlight the need to directly and simultaneously compare encoding across the SC-FEF-LIP network to determine their contributions to computing abstract category information from sensory representations in upstream visual cortical regions (e.g. MT and MST) [39, 51, 52].

Monkeys were trained on the DMC task for hundreds of training sessions, spanning a period of many months, prior to neuronal recordings. It will be interesting to investigate whether abstract encoding in subcortical or motor structures such as the SC emerges only after prolonged training on a task. Task-related encoding might emerge at different rates or at different learning stages in the LIP and the SC; the SC may be recruited to participate in a cognitive task only once it is highly familiar. Alternatively, the SC might participate more broadly in learning and performing complex tasks and behaviors than previously appreciated, even during early stages of learning a task.

Our findings are interesting to consider from an evolutionary perspective, and highlight the importance of considering the functions of the SC between mammals and other vertebrates. In order to respond appropriately to stimuli, animals must rapidly combine sensory encoding with more abstract, learned knowledge. While previous work in mammalian SC has emphasized its role in simple sensory-motor mapping, our work raises the prospect SC also mediates more flexible, and even cognitive, behaviors. This could be mediated by learning-dependent plasticity within SC and/or contextual/cognitive inputs from higher brain centers. Indeed, it could be advantageous for an area like the SC, which is close to both sensory input and motor output brain centers, to play such a role in order to facilitate rapid yet flexible behaviors. The idea that the SC is involved in mediating complex behaviors is especially plausible in non-mammalian vertebrate species that lack a neocortex, in which the tectum occupies a much larger fraction of brain volume and is known to play a major role in visual processing. Indeed, studies have found innate spatial encoding of stimulus size category in the optic tectum of untrained barn owls [53, 54]. In mammals and primates, this spatial orienting circuit may have evolved to rapidly compute more complex types of information (like the visual categories described here), while cortical pathways developed to allow for slower, but even more sophisticated processing.

## Methods

### Subjects

Two adult (13-15 years old) male rhesus macaques (Macaca mulatta) participated in the experiment (Monkey N: ~12 kg, Monkey S: ~13 kg). All procedures were in accordance with the University of Chicago Institutional Animal Care and Use Committee and the National Institutes of Health guidelines and policies.

### Behavioral tasks

For the behavioral tasks described below, the monkeys were head restrained and seated in a primate chair inserted inside an isolation box (Crist Instrument), facing a 24-inch LCD monitor on which stimuli were presented (1,920 × 1,080 resolution, refresh rate 60 Hz, 57 cm viewing distance). Reward delivery, stimulus presentation, behavioral signals, and task events were controlled by MonkeyLogic software [55], running under MATLAB on a Windows-based PC. Gaze position was measured with an optical eye tracker (Eyelink 1000; SR Research, Ottowa, Canada) with a 1.0 kHz sample rate. For both tasks, monkeys initiated trials by holding a manual touch bar.

#### Delayed match-to-category task

We trained monkeys to perform a delayed match-to-category (DMC) task in which they grouped twelve dot-motion stimuli into two categories based on two orthogonal boundaries, such that motion directions that are 180° apart belong to the same category. Motion directions were separated into quadrants with three directions per quadrant, and stimuli within the same quadrant were 22.5° apart and near-boundary directions were 22.5° away from the boundary. The stimuli were 6°-diameter circular patches of white dots moving at a speed of 10°/s with 100% coherence, presented at 6.5-7.5° eccentricity in the contralateral visual field. Animals were required to fixate within a 2.5-3.5° window.

#### Memory-guided saccade task

We used a memory-guided saccade (MGS) task to identify visual and motor receptive fields of LIP and SC neurons (**Extended Fig. 7**). At the start of a trial, monkeys had to maintain fixation on the central cue for 500 ms, after which a white square target briefly appeared for 300 ms at one of eight peripheral locations (equally spaced and concentric at 6.5° eccentricity; see **Extended Fig. 7b**). The target presentation was followed by a 1000-ms delay period, after which the fixation cue disappeared and monkeys had to saccade to the remembered location of the visual target presented earlier in the trial.

### Surgical procedures and electrophysiological recordings

Monkeys were implanted with a titanium headpost and a single recording chamber positioned over LIP and SC. Stereotaxic coordinates for chamber placement were determined from magnetic resonance imaging (MRI) scans obtained before implantation of recording chambers. LIP and SC recordings were conducted in separate sessions, typically using 16- and 24-channel linear Plexon V-probes (in which channels span 1.5-2.0 mm of tissue), a dura-piercing guide tube, and a NAN microdrive system (NAN Instruments). A small subset of recording sessions from one monkey were conducted using single epoxy-insulated tungsten electrodes (FHC, Inc). We used anatomical landmarks and responses during the MGS task to guide recordings. For SC recordings, we primarily targeted neurons in superficial and intermediate layers, although we also recorded neurons in deep layers as well due to the ~2mm span of recording channels on our probes. Neurophysiological signals were amplified, digitized, and stored for offline spike sorting (Plexon) to verify the quality and stability of neuronal isolation.

### SC inactivation

We infused muscimol, a GABA_A_ agonist, to unilaterally inactivate the SC. We built a microfluidic injectrode system to deliver small amounts of the drug or saline (0.25-0.5 μL; see **Extended Table 1**) using the protocol developed by [56]. To ensure that we precisely injected the drug into superficial and intermediate layers of the SC, we used a custom 16-channel Plexon S-probe with a fluid delivery channel that allowed us to monitor neural activity during probe lowering and before injection. Before drug injection on each session, monkeys first completed a control behavioral session in which they performed at least 200 correct trials of the DMC task and at least 100 correct trials of the MGS task (Monkey N) or visually-guided saccade task (Monkey S). After monkeys completed the control behavioral session, we infused the drug and waited 15-25 minutes to begin the post-treatment behavioral session. To verify success of SC inactivation, we compared saccade metrics (peak saccade velocity) during the MGS/VGS tasks during the control and post-treatment trials. We analyzed data from 12 muscimol injections sessions (Monkey N: 6 sessions, Monkey S: 6 sessions) and six control saline injection sessions (Monkey N: 2 sessions, Monkey S: 4 sessions). **Extended Table 1** provides information for each injection session, including muscimol and saline concentration, injection volume, and number of completed DMC trials.

### Behavioral inclusion criteria

For electrophysiological recordings and inactivation experiments, we included sessions in which behavioral performance on each category for the DMC task was at least 75% (criterion applied only to control blocks for the inactivation experiments). We excluded six LIP recording sessions (four in Monkey N and two in Monkey S) from analyses due to poor behavioral performance. For the inactivation experiments, we excluded two sessions in Monkey S (one saline injection session with 66% accuracy for category 2 during the control block, and one muscimol injection session with 53% accuracy for category 2 during the control block).

For DMC analyses, we included well-isolated neurons for which we had data recorded during at least five correct trials for each sample direction. We analyzed spiking data during the DMC task from 555 LIP neurons recorded over 49 recording sessions (Monkey N: *N* neurons = 228, *N* sessions = 36; Monkey S: *N* neurons = 327, *N* sessions = 13) and 604 SC neurons recorded over 38 recording sessions (Monkey N: *N* neurons = 362, *N* sessions = 26; Monkey S: *N* neurons = 242, *N* sessions = 12). We collected and analyzed spiking activity during the MGS task in a subset of 423 SC neurons (Monkey N: *N* = 259, Monkey S: *N* = 164) for which we recorded data from least two correct trials for each MGS condition.

### Data analysis

All analyses were performed in Python v3.7.3. Behavioral analyses for the DMC task (including those for inactivation) were performed on all completed trials (i.e., correct trials, misses on match trials, and false alarms on non-match trials). Unless otherwise specified, all neural analyses for the DMC task were performed only on correct trials. Behavioral and neural analyses for the MGS/VGS tasks were performed only on correct (completed) trials. All *P*-values are two-tailed unless otherwise specified. For neural analyses, spike trains for each neuron were smoothed using Gaussian kernel (σ = 30 ms). Eye tracker gaze position data were low-pass filtered to reduce noise using a 2nd-order Butterworth filter with a 70-Hz cutoff.

#### Behavioral performance

To compare differences in mean behavioral accuracy (across all sample directions) between LIP and SC recording sessions in each monkey (**Extended Fig. 1a**), we used a permutation test in which we randomly permuted mean accuracy values between the two brain areas (while preserving the number of sessions per area). We repeated this procedure for 5000 unique iterations to generate a null distribution of accuracy differences. To compare differences in behavioral performance on match trials in which the sample and test stimuli were in the same vs. opposite quadrants, we computed the difference in mean accuracy for same vs. opposite quadrant match trials for each session and used a permutation test (with 5000 iterations) to randomly permute the per-session accuracy values between the two conditions.

#### Quantifying single-neuron category tuning

We quantified the strength, reliability, and time course of single-neuron category tuning using a receiver operating characteristic (ROC)-based category tuning index (rCTI)[41]. For each neuron, we applied ROC analysis to distributions of trial-by-trial firing rates and compared area under the ROC curve (AUC) values for eight pairs of sample motion directions that are in the same category (Within-Category; WC) and eight pairs of directions that are in different categories (Between-Category; BC). To ensure equalized angle differences between WC and BC pairs (and thus prevent direction tuning from contaminating rCTI), the WC and BC groups each included four direction pairs spaced 45° apart and four direction pairs spaced 135° apart (**Extended Fig. 2a**). We quantified rCTI at each timepoint as the mean rectified WC AUC subtracted from the mean rectified BC AUC:

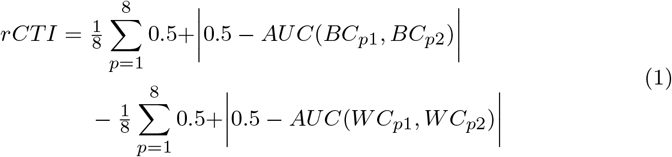

where *BC_p1_* and *BC_p2_* are the two directions in the *p^th^* BC pair, and *WC_p1_* and *WC_p2_* are the two directions in the *p^th^* WC pair.

We applied the rCTI analysis to smooothed spike trains (see above) across 5-ms time steps in the trial. To generate the error shading shown in **Fig. 2c** and **d**, we calculated rCTI for each neuron over 500 bootstraps using 15 trials per sample motion direction (with replacement). We generated null distributions of rCTI values for each neuron using a bootstrap analysis (repeated 5000 times) in which we randomly assigned (with replacement) eight direction pairs (four 45°-spaced and four 135°-spaced pairs) to each of the shuffled BC and WC groups (**Extended Fig. 2**). We defined category-tuned “runs” as time bins at which rCTI values significantly exceed the null distribution for a minimum of six consecutive analysis time bins (30 ms). We considered neurons to be category-tuned if they had at least one significant run, and defined latency of category selectivity for each category-tuned neuron as the first time bin of the earliest significant run.

To test for significant above-chance mean rCTI in each brain area (as shown in **Fig. 3f**), we used a permutation procedure in which we computed a null mean rCTI across neurons for each WC/BC-label-shuffling iteration. To test for a significant difference between brain areas in the onset time of category selectivity for mean rCTI, we compared the observed between-area latency difference to a null distribution of latency differences. For each of 5000 iterations, we randomly permuted neurons between the two areas (while preserving the number of neurons in each area) and we computed the difference in latency of category selectivity onset in the two shuffled groups. To test for differences in onset time of category selectivity between Vis and Vis-other SC neurons, we used a similar procedure in which we randomly permuted neurons between the two groups instead of between areas.

#### Support vector machine (SVM) analyses

We used SVM classifiers (with a linear kernel) to quantify the strength and timing of sample stimulus category encoding in populations of LIP and SC neurons. To quantify category encoding in a direction-independent manner, we constructed cross-quadrant classifiers for which training sets consisted of trials in which the sample motion directions were from two of the four quadrants (one from each category), and testing sets consisted of sample motion direction trials from the other two quadrants (**Fig. 2g**). The training and testing quadrants were randomly chosen on each iteration. The analysis was applied in 5-ms steps across time in the trial and repeated for 200 iterations. For each neuron, we included 15 trials from each of six sample motion directions for training (as described above) and 15 trials from each of the remaining six sample motion directions for testing. To reduce the biases in classifier performance across brain areas due to an unequal number of neurons, for each iteration of the analysis, we randomly selected *N* neurons for inclusion, where *N* is the number of neurons in the brain area with the lower number of neurons. We generated null distributions of decoder performance values at each time using a permutation procedure (repeated 1000 times) in which we shuffled the sample direction label assigned to each trial.

We also used linear SVM classifiers to decode sample direction from LIP and SC population activity. To quantify the amount of direction encoding in a category-independent manner, the training/validation sets for each iteration of the classifier only included data from one of the two categories. The classifiers were trained on 48 trials (8 trials from each of the six directions from one of the two categories, randomly chosen) and validated on 12 held-out trials (2 trials from each of the six motion directions). This analysis was applied in 5-ms steps across the trial and repeated for 200 iterations.

#### Identifying task-responsive neurons

To identify neurons that are task-responsive during the DMC task, we used a bin- and parameter-free statistical test to detect any consistent time-locked modulations in firing rate for each neuron [57]. In brief, this analysis consists of the following steps (applied separately for each sample direction): (1) aligning the spike trains for all correct trials to the onset of the sample stimulus, (2) stacking these spike train to create a single vector of spikes relative to sample onset, (3) calculating the cumulative distribution of spikes over trial time using this spike vector, and (4) comparing this cumulative distribution to a linear baseline (which represents an unvarying firing rate over time), producing a deviation value for each timepoint. To generate a null distribution of 5000 deviation-from-baseline values, we shuffled the spike trains in each trial to destroy any time-locked activity patterns across trials while preserving the total number of spikes, and computed the maximum deviation (across time) for these shuffled data. For this analysis, we included data for each trial from 500 ms before sample stimulus onset (i.e., the start of the pre-sample fixation period) until the end of the first test stimulus epoch. We also computed the peak mean firing across time (in the period from the beginning of the sample epoch until the end of the first test epoch) for each sample direction. We classified each neuron as task-responsive if it satisfied the following two criteria: (1) if it showed a significant modulation in firing rate (i.e., had significantly elevated deviation-from-linear-baseline values) at any timepoint from the start of the sample epoch until the end of the test epoch, and (2) if its maximum peak mean firing rate (across sample directions) was at least 3 sp/s. In LIP, 506/555 (91.2%) of neurons were task-responsive (Monkey N: 210/228, 92.1%; Monkey S: 296/327, 90.5%), and in SC, 504/604 (83.4%) of neurons were task-responsive (Monkey N: 300/362, 82.9%; Monkey S: 204/242, 84.3%).

#### Identifying direction-tuned neurons during the DMC task

To identify neurons that are significantly direction tuned during the sample epoch of the DMC task, we computed a direction tuning index (DTI) for each neuron using the circular variance method introduced in [58]. We calculated neurons’ mean firing rate for each sample stimulus direction in a direction vector space, and quantified DTI as the normalized length of the sum of these vectors:

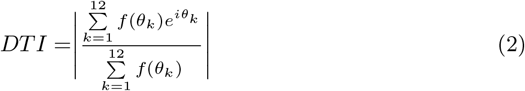

where f(*θ*_k_) is a neuron’s mean firing rate for direction *θ*_k_.

To test for significant direction tuning, we compared the true DTI to a distribution of 5000 null DTIs generated by randomly shuffling the direction labels assigned to each mean firing rate. We applied this analysis to firing rates in three 200-ms windows from 0 to +600 ms relative to sample stimulus onset. We classified neurons as direction tuned if they showed significant direction tuning during at least one of the three time windows and if they were identified as responsive during the DMC task (see above). In LIP, 153/506 (30.2%) of neurons were significantly direction-tuned (Monkey N: 79/210, 39.5%; Monkey S: 74/296, 25.0%), and in SC, 103/504 (20.4%) of neurons were significantly direction-tuned (Monkey N: 53/300, 17.7%; Monkey S: 50/204, 24.5%).

### Memory-guided saccade task analyses

#### Identifying visually-responsive neurons

We analyzed neuronal activity during the MGS task in order to characterize the visual and motor response fields of SC neurons. For each neuron, we determined whether the DMC stimulus was presented in its visual receptive field by identifying the MGS condition (MGS_DMCloc_) whose location overlapped with the DMC stimulus location on that session. We then determined whether the neuron was significantly modulated during the MGS visual epoch for that condition. For each of the eight MGS conditions, we computed the mean firing rate (per trial) in 25-ms windows during the visual epoch (from 0 to +400 ms relative to stimulus onset). We used the Kruskal–Wallis H-test to compare the firing rate distributions across these windows, and compared the resulting H-statistic to a distribution of 5000 null H-statistics. To generate the null H distribution, we shuffled the neuron’s time-varying firing rates (from 0 to +400 ms relative to stimulus onset) for each trial, calculated the mean firing in 25-ms windows for these permuted trials, and computed a shuffled H-statistic. Neurons were classified as “Vis” neurons if they was significantly modulated across the visual stimulus period for the MGS_DMCloc_ condition and if their maximum firing rate across analysis windows and conditions was above 3 sp/s. Neurons were classified as “Vis-other” if they were significantly modulated across the visual stimulus period for another MGS condition (and if their maximum firing rate was above 3 sp/s). Neurons were classified as “Non-vis” if they were not significantly modulated across the visual period for any of the MGS conditions, or if their maximum firing rate across analysis windows and conditions was below 3 sp/s. 115 (27.2%) neurons were classified as Vis neurons (Monkey N: 82 [31.7%], Monkey S: 33 [20.1%]), 178 (42.1%) as Vis-other neurons (Monkey N: 97 [37.5%], Monkey S: 81 [49.4%]), and 130 (30.7%) as non-Vis neurons (Monkey N: 80 [30.9%], Monkey S: 50 [30.5%]).

#### Identifying saccade-modulated neurons

We analyzed activity of SC neurons during the saccade period of the MGS task (−200 ms to +50 ms relative to saccade onset) to identify neurons that are significantly modulated by saccade direction. For each neuron, we computed its mean firing rate across the saccade period window for each trial. We used the Kruskal–Wallis H-test to compare the neuron’s the firing rate distributions across the eight MGS conditions, and compared the resulting H-statistic to a distribution of 5000 null H-statistics generated by shuffling condition labels among trials. 145/423 (34.3%) neurons were significantly modulated by saccade direction (Monkey N: 96/259 [37.1%], Monkey S: 49/164 [29.9%]).

#### Task modulation index

To quantify differences in modulation during the MGS and DMC tasks for each SC neuron, we computed a task-preference index (**Fig. 3b**). We defined amount of MGS modulation as the range of mean firing rates across conditions during the peri-saccade period of the MGS task (−200 to +50 relative to saccade onset), and the amount of DMC modulation as the range of mean firing rates across sample directions during the sample epoch of the DMC task (+150 tp + 550ms relative to stimulus onset).

We then normalized these ranges for each neuron by the neuron’s overall firing rates across all times/conditions/tasks. The task-preference index was defined as the ratio of DMC modulation to MGS modulation. We used a Hartigan’s dip test to test for bimodality in the distribution of task-preference indices across all SC neurons.

### Linear encoding models

We constructed linear encoding models [45] to quantify how much firing rates of individual neurons (across trials and time within trials) are modulated by stimulus category vs microsaccades. The linear models contained regressors related to stimulus category and microsaccade parameters. For the category regressors, we constructed a binary vector containing a pulse at the time of the sample stimulus onset, and created copies of this vector shifted in time by 1ms for every point until the end of the trial. The microsaccade regressors included two analog regressors: horizontal and vertical eye velocity at each timepoint throughout the trial, shifted in time by −50 relative to neural activity to account for lag between neural activity and saccades. We also included two types saccade event kernel regressors: (1) a binary vector containing a pulse at every timepoint at which a microsaccade occurred, and (2) a vector containing microsaccade direction at every timepoint at which a microsaccade occurred and zeros at every other timepoint. We created time-shifted copies of the binary saccade vector and saccade direction vector, spanning from −500 until +100 ms (relative to saccade onset) in 10-ms steps. The design matrix of the full model included all of the category and saccade regressors. We also built reduced models that contained shuffled saccade regressors and unshuffled category regressors, or shuffled category regressors and unshuffled saccade regressors. For each neuron, we fit the models using ridge regression (with L2 regularization and 10-fold cross validation) and computed an R^2^ for the full model and each of the reduced models. To quantify how well category or saccade regressors predict neural activity in each neuron, we computed the change in cross-validated R^2^ from the full model to each reduced model. A large (negative) change in R^2^ indicates a strong contribution of the excluded variables.

### Subspace alignment analysis

We used a subspace alignment analysis introduced in [46] to quantify the degree of alignment between neural activity in the SC during the MGS and DMC tasks. For this analysis, we constructed matrices *D* and *M* of neural activity during the DMC and MGS tasks, respectively. *D* and *M* were size *N* by *cxt*, where *N* is the number of neurons, c the number of conditions (12 for DMC and 8 for MGS), and *t* is the number of time points per condition. Each row of *D* and *M* contains the concatenated mean firing rates (per condition and across time points) of one neuron. We normalized the firing rates of each neuron by its range (across all included DMC and MGS conditions and time points) plus a constant, chosen as 10 sp/s. We then performed principal components analysis (PCA) on the matrix *D* to obtain the top 12 DMC PCs, and on matrix *M* to obtain the top 12 MGS PCs. We then projected the DMC activity *D* onto both the DMC and MGS PCs and calculated sum of the percent of variance explained (relative to total variance of *D*) for each of the projections. We quantified the alignment index (AI) between the two subspaces as the ratio of these two sums. The logic behind this analysis is that if the DMC and MGS subspaces are approximately orthogonal, the projection of *D* onto the MGS PCs will capture minimal *D* variance. AI ranges from 0 (indicating perfect orthogonality between two subspaces) and 1 (indicating perfect alignment).

To determine whether measured AI values are more (or less) misaligned than expected by chance, we calculated the alignment between pairs of random subspaces sampled from the full covariance structure of the data to generate a null distribution of alignment values [46]. To create the random subspaces, we first computed the covariance matrix *C* from the concatenated *D* and *M* matrices, and obtained the left singular vectors (*U*) and singular values (*s*) of *C* using singular value decomposition. For each of 5000 iterations per comparison, we computed the AI between two random subspaces (*v_rand_*). We sampled each random subspace *v_rand_* as follows:

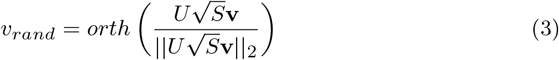

where **v** is an *N* × 12 matrix in which each element is drawn from a normal distribution with mean 0 and variance 1, and *orth*(*X*) returns the orthonormal basis of X defined by its left singular values.

For the main alignment analysis (shown in **Fig. 3c-d**), we included DMC task data from 150-550 ms after sample stimulus onset (during stimulus presentation) and MGS task data from 200 before saccade onset until saccade onset. Data for both tasks were sampled in 10-ms steps. Note that results were equivalent when we used time windows of equal length for the two tasks (MGS: −200 ms to +0 ms relative to saccade onset, DMC: +350 to +550 ms relative to sample onset; Monkey N: AI = .134, *P* < .001, Money S: AI = .233, *P* < .001)).

### SC inactivation analyses

To verify efficacy of SC inactivation, we quantified the difference in peak saccade velocity for saccades towards the inactivated hemifield during the MGS/VGS task between the control and post-treatment blocks. For each trial, we computed the maximum eye gaze velocity from 200 ms before go cue onset until successful target fixation initiation. We excluded one trial for Monkey N (session 2, muscimol treatment, upper-center condition) in which we could not accurately quantify peak saccade velocity because the monkey blinked during the response period. For each session, we combined trials from the three conditions in which the target was in the inactivated hemifield (“Contralateral”), and the three conditions in which the target was out of the inactivated hemifield (“Ipsilateral”). We tested for significant differences in mean peak saccade velocity between the control and treatment blocks for Contralateral and Ipsilateral trials on each session using a bootstrap test with 5000 iterations (**Extended Table 2**). We also tested for significant differences in mean peak saccade velocity between control and treatment blocks for Contralateral trials pooled across all muscimol sessions and pooled across all saline sessions, as shown in **Fig. 4d**. For the muscimol sessions, this analysis included 270 (470) control (treatment) trials in Monkey N and 320 (376) control (treatment) trials in Monkey S, and for saline sessions included 72 (103) control (treatment) trials in Monkey N and 207 (215) control (treatment) trials in Monkey S.

We used a two-sided Fisher exact test to quantify differences in behavioral performance on the DMC task between control and post-treatment blocks for each session. The statistics for the test are shown in **Extended Table 3**.

## Data and Code Availability

All data and code used to generate the figures and results in this manuscript will be made publicly available prior to publication via Figshare (figshare.com).

## Acknowledgments

We are grateful to the following individuals for helpful discussions and/or comments on an earlier version of this manuscript: Chris Hauser, Matthew Kaufman, Kenneth Latimer, Jason MacLean, John Maunsell, Krithika Mohan, Matthew Rosen, Vinay Shirhatti, and Yang Zhou. We are grateful for expert technical assistance from Suha Chang, Yinghui Qiu, and Sam Zheng. We acknowledge expert animal care and veterinary support from the staff the University of Chicago Animal Resources Center. This work was supported by NIH R01EY019041, NIH U19NS107609, NIH NRSA 1F31MH124395 (BP), NIH NRSA F30EY033648 (OZ), NIH NRSA F31 EY029155 (WJJ), and DOD Vannevar Bush Faculty Fellowship N000141912001 (DJF).

## Extended figures and tables

**Extended Fig. 1:**
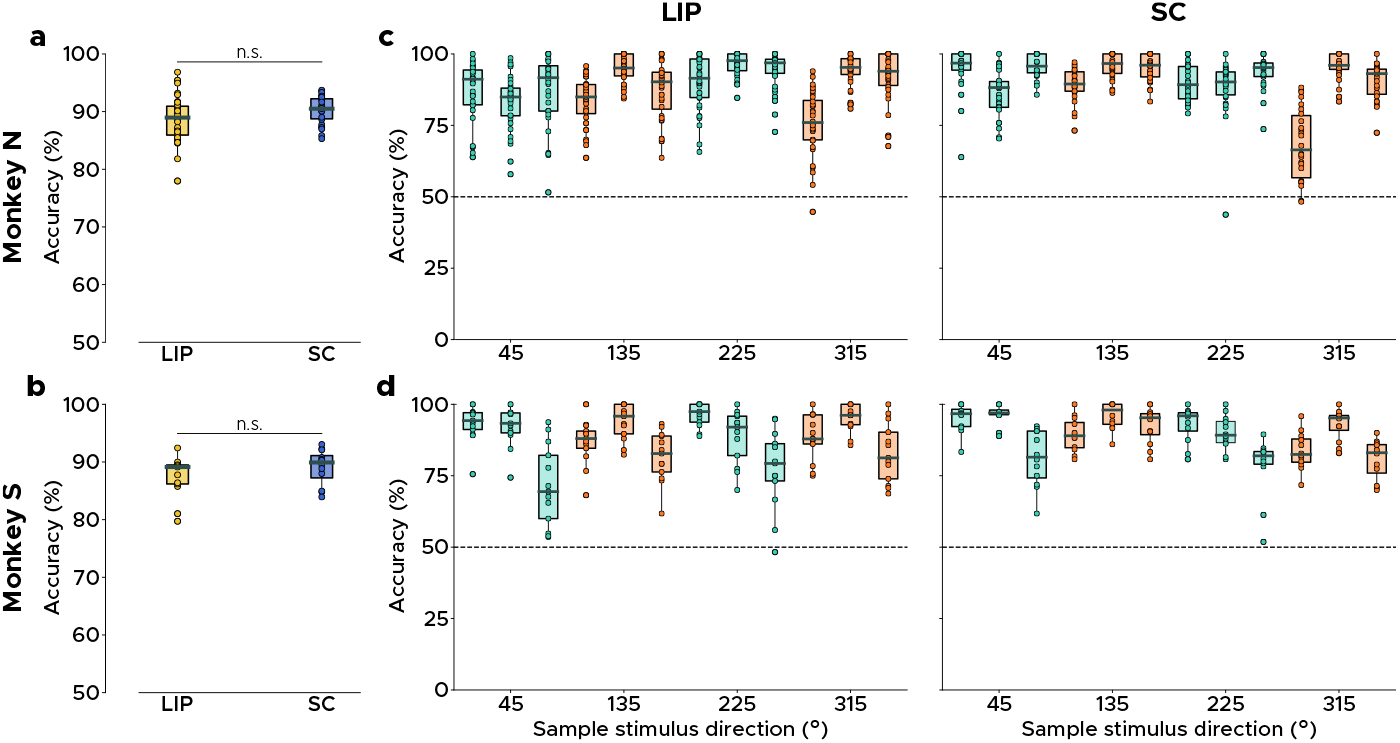
No difference in behavioral performance between LIP and SC recording sessions. **a,** Distributions of overall session accuracy for LIP and SC recording sessions in Monkey N. There was no difference in mean accuracy between areas (LIP: 88.8 ± 3.9%, SC: 90.3 ± 2.3%, *P* = .094, permutation test). **b,** same as **a** but for Monkey S (LIP: 87.4 ± 3.5%, SC: 89.1 ± 2.9%, *P* = .229, permutation test). **c,** Mean accuracy by sample direction for LIP (left) and SC (right) recording sessions. **d,** same as c but for Monkey S.

**Extended Fig. 2:**
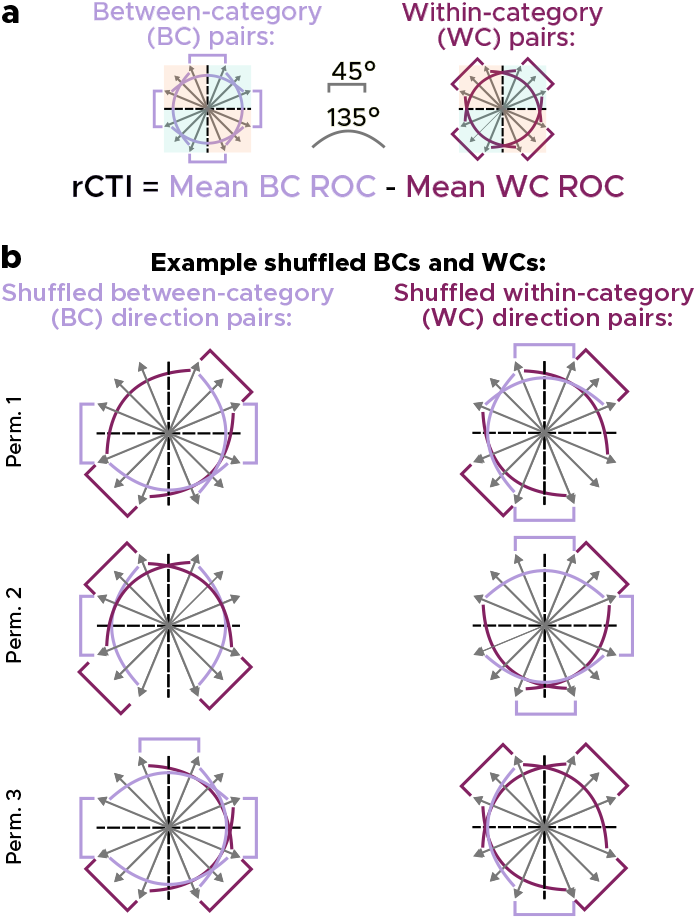
rCTI method and shuffling procedure. **a,** Schematic of the rCTI analysis used to quantify strength of category tuning in each neuron. rCTI compares ROC values between pairs of sample directions that are in the same category (Within-Category; WC) vs different categories (Between-Category; BC). WC and BC groups each consisted of eight sample directions (four pairs spaced 45° apart and four pairs spaced 135° apart. **b,** We generated null distributions of rCTI values across timepoints for each neuron using a shuffling procedure in which we reshuffled the labels (between- vs within-category) assigned to each pair of directions, such that each shuffled group contained four 45°-apart direction pairs and four 135°-apart direction pairs. The permutation procedure was repeated 4900 times, once for every combination of shuffled direction pairs that conformed to the criterion above. Bracket colors denote true group assignment of a direction pair (light purple = between-category, dark purple = within-category).

**Extended Fig. 3:**
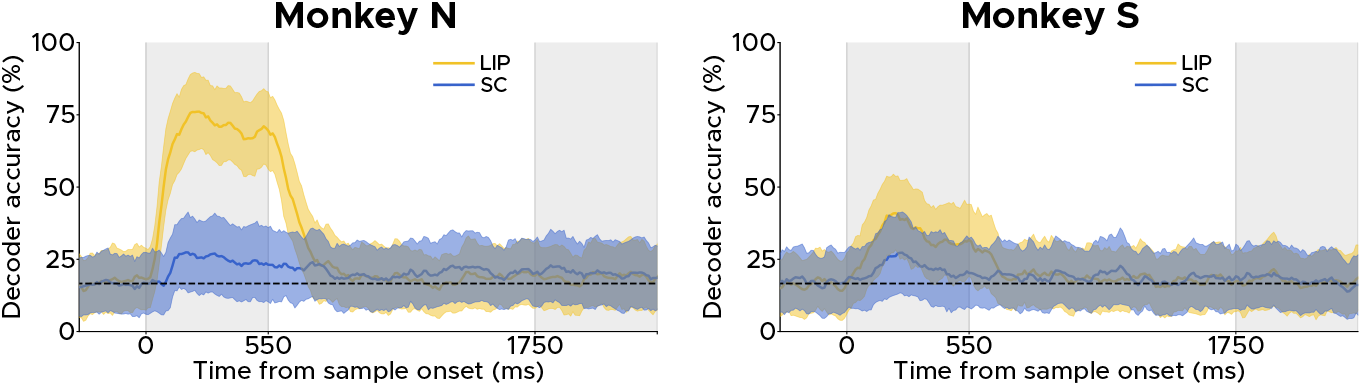
Direction classifier accuracy in LIP and SC. Time course of categoryindependent direction classifier accuracy across LIP and SC neurons. Lines and shading indicate mean ± s.d.

**Extended Fig. 4:**
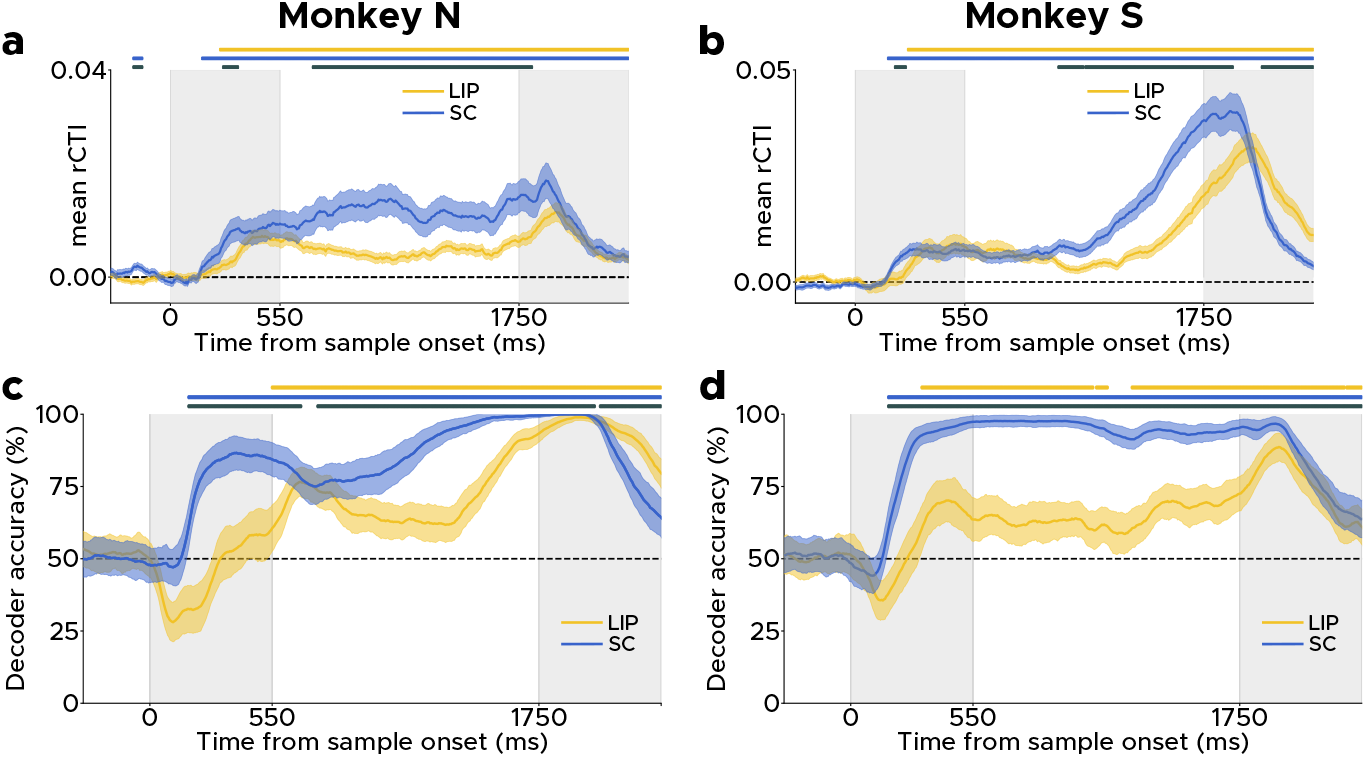
rCTI and category classifier accuracy in LIP and SC by monkey. **a,** Mean Time course of rCTI across LIP and SC neurons in Monkey N. Lines and shading indicate mean ± s.d. rCTI. **b,** same as a but for Monkey S. **c,** Time course of cross-quadrant category decoding accuracy in LIP and SC for Monkey N. Lines and shading indicate mean ± s.d. decoding accuracy. **d,** same as c but for Monkey S. In all panels, yellow (blue) symbols above plot indicate timepoints at which LIP (SC) values are significantly above chance, and black symbols indicate time points at which there is a significant difference between LIP and SC values when either or both area are significantly above chance (permutation test, all *P* < .050).

**Extended Fig. 5:**
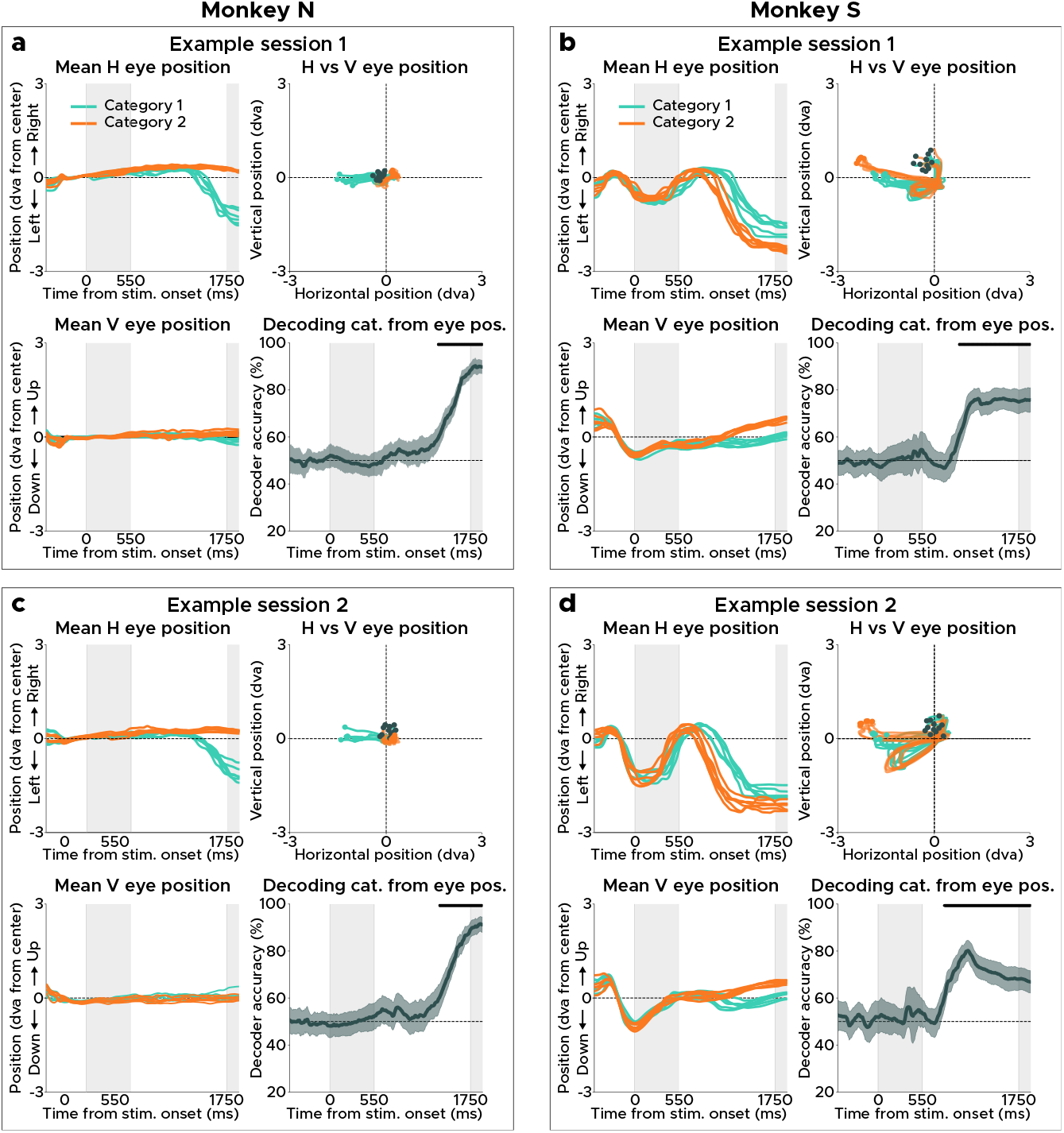
Monkeys’ eye movements reflect working memory contents during delay period of DMC task. **a,** Top left: Mean horizontal eye position across trial time for each sample motion direction during an example session. Bottom left: Mean vertical eye position across trial time. Top right: Mean horizontal vs vertical eye position. Black circles indicate mean eye position at the begining of the trial, and colored circles indicate the mean eye position at the beginning of the delay period. Bottom right: Accuracy of cross-quadrant category classifier trained on eye position. Black symbols above panel indicate time points at which the category classifier performs significantly above chance (*P* <0.05, permutation test). Shading indicate s.d. **b,** Same as a but for Monkey S. **c, d** Same as a and b but for another example session.

**Extended Fig. 6:**
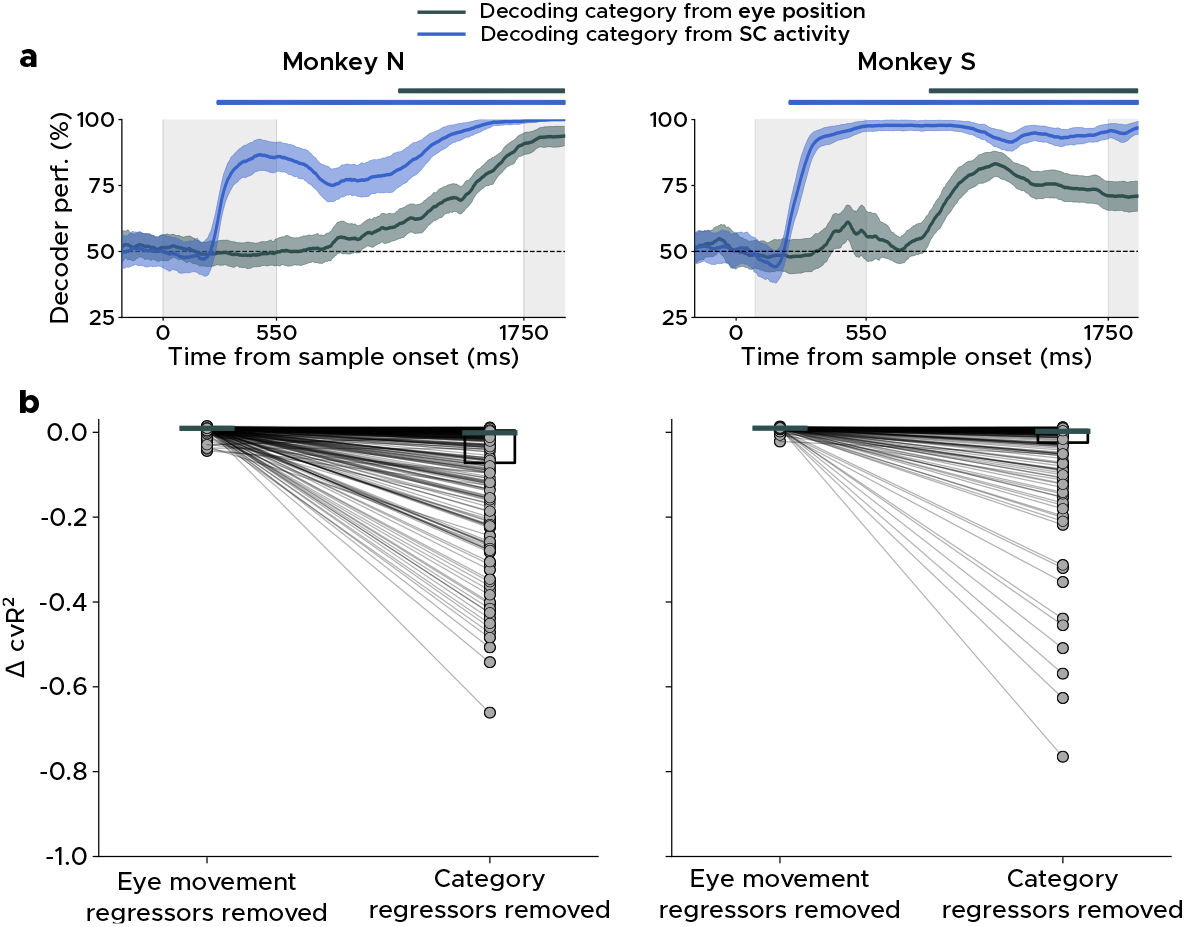
Comparison of the contribution of stimulus category and eye movements to single-neuron activity. **a,** Time course performance of category classifiers trained on SC neural data (blue) or on horizontal and vertical eye gaze trajectories across all SC recording sessions (grey). Symbols above panels indicate time points at which classifier accuracy performed significantly above chance (*P* <0.05, permutation test). **b,** Change in cross-validated R^2^ values from full linear encoding models (which include both eye movements and category regressors) to reduced linear models that with either eye movement-related or category regressors removed.

**Extended Fig. 7:**
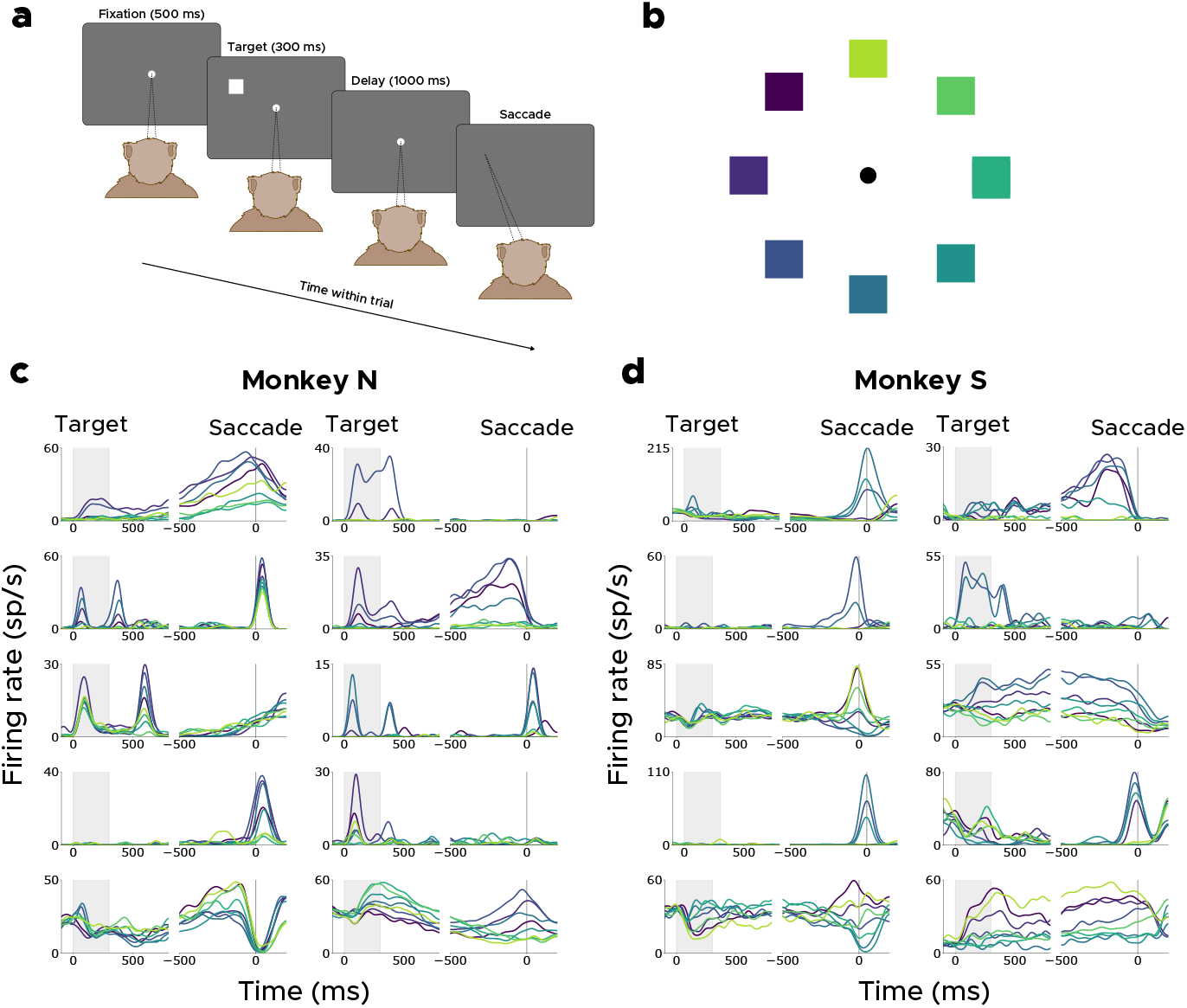
SC responses during the memory-guided saccade (MGS) task. **a,** Schematic of the MGS task. **b,** Color-coding scheme of the eight stimulus positions used in the MGS task. **c,** Example PSTHs of SC neurons during the MGS task in Monkey N. Each trace corresponds to one of the right stimulus positions illustrated in **b**. **d,** same as c but for Monkey S.

**Extended Fig. 8:**
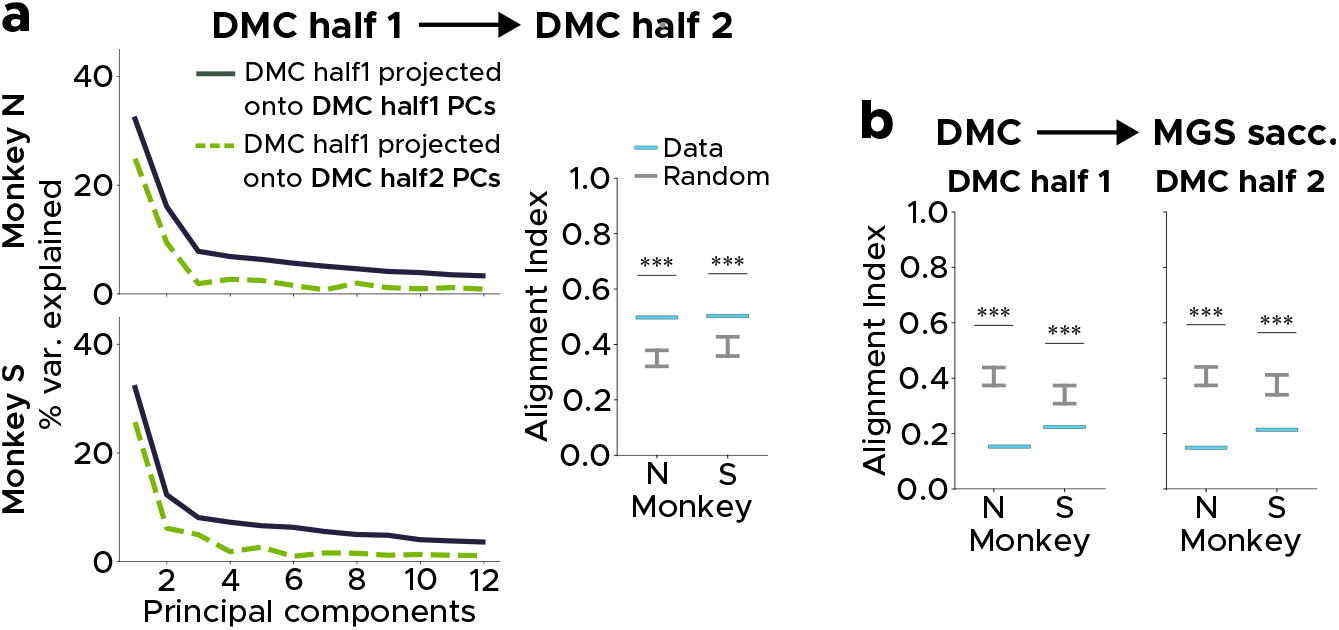
Additional subspace alignment indices. **a,** Alignment between SC neural subspaces recorded during the first half and second half of the trials for each session. Data are significantly more aligned than chance (Monkey N: AI = .499, *P* < .001, Money S: AI = .505, *P* < .001). **b,** Left: SC neural subspaces during the first half of DMC trials and the MGS saccade period were significantly misaligned (Monkey N: AI = .155, *P* < .001, Money S: AI = .225, *P* < .001). Right: SC neural subspaces during the second half of DMC trials and the MGS saccade period were significantly misaligned (Monkey N: AI = .151, *P* < .001, Money S: AI = .216, *P* < .001).

**Extended Table 1:** SC inactivation session information

| Monkey | Exp | Treatment | Concentr.<br>($\mu\text{g}/\mu\text{L}$) | Injection vol.<br>($\mu\text{L}$) | # DMC trials<br>(control) | # DMC trials<br>(injection) |
| --- | --- | --- | --- | --- | --- | --- |
| N | 1 | Saline | 8 | 0.50 | 288 | 292 |
| N | 2 | Muscimol | 5 | 0.33 | 308 | 222 |
| N | 3 | Muscimol | 5 | 0.33 | 275 | 353 |
| N | 4 | Muscimol | 5 | 0.33 | 251 | 510 |
| N | 5 | Muscimol | 5 | 0.25 | 331 | 456 |
| N | 6 | Muscimol | 5 | 0.25 | 294 | 385 |
| N | 7 | Muscimol | 5 | 0.25 | 359 | 218 |
| N | 8 | Saline | 8 | 0.25 | 273 | 369 |
| S | 1 | Saline | 8 | 0.25 | 285 | 427 |
| S | 2 | Muscimol | 5 | 0.25 | 302 | 278 |
| S | 3 | Muscimol | 5 | 0.25 | 403 | 104 |
| S | 4 | Muscimol | 5 | 0.25 | 217 | 138 |
| S | 5 | Saline | 8 | 0.25 | 258 | 285 |
| S | 6 | Saline | 8 | 0.25 | 281 | 547 |
| S | 7 | Muscimol | 5 | 0.25 | 258 | 164 |
| S | 8 | Muscimol | 5 | 0.25 | 282 | 182 |
| S | 9 | Saline | 8 | 0.25 | 305 | 312 |
| S | 10 | Muscimol | 5 | 0.25 | 290 | 240 |

**Details of injection experiments in two monkeys.** Each row contains an individual experimental session, and each column contains the details of that session. The column descriptions are as follows: “Monkey”: indicates which monkey the experiment was performed with; “Exp”: session number by animal; “Treatment”: indicates whether the injection was saline or muscimol; “Concentr.”: concentration of muscimol or saline (in μg/μL); “Injection vol.”: total injection volume (in μL); “N DMC trials (control)”: total number of completed trials (correct and incorrect) for the DMC task during the *control* (pre-injection) block; “N DMC trials (injection)”: total number of completed trials (correct and incorrect) for the DMC task during the *post-injection* block.

**Extended Table 2:** MGS peak saccade velocity (PSV) during SC inactivation experiments

| Monkey | Exp | Contralateral PSV (°/s) |  |  | Ipsilateral PSV (°/s) |  |  |
| --- | --- | --- | --- | --- | --- | --- | --- |
| | | Ctrl<br>($\mu \pm \sigma$) | Inject<br>($\mu \pm \sigma$) | P-val | Ctrl<br>($\mu \pm \sigma$) | Inject<br>($\mu \pm \sigma$) | P-val |
| N | 1<sup>†</sup> | 434 ± 105 | 459 ± 96 | .318 | 314 ± 75 | 300 ± 64 | .370 |
| N | 2 | 387 ± 96 | 232 ± 47 | <.001* | 282 ± 45 | 334 ± 45 | <.001<sup>⊗</sup> |
| N | 3 | 381 ± 68 | 237 ± 38 | <.001* | 273 ± 43 | 329 ± 44 | <.001<sup>⊗</sup> |
| N | 4 | 447 ± 108 | 243 ± 51 | <.001* | 260 ± 51 | 302 ± 46 | <.001<sup>⊗</sup> |
| N | 5 | 432 ± 94 | 257 ± 49 | <.001* | 269 ± 54 | 307 ± 62 | <.001<sup>⊗</sup> |
| N | 6 | 443 ± 79 | 264 ± 52 | <.001* | 264 ± 46 | 297 ± 55 | <.001<sup>⊗</sup> |
| N | 7 | 416 ± 62 | 270 ± 57 | <.001* | 245 ± 59 | 310 ± 57 | <.001<sup>⊗</sup> |
| N | 8<sup>†</sup> | 431 ± 79 | 421 ± 90 | .579 | 252 ± 67 | 264 ± 60 | .344 |
| S | 1<sup>†</sup> | 273 ± 51 | 292 ± 65 | .123 | 287 ± 65 | 299 ± 76 | .317 |
| S | 2 | 260 ± 111 | 193 ± 84 | <.001* | 452 ± 55 | 592 ± 108 | <.001<sup>⊗</sup> |
| S | 3 | 430 ± 67 | 174 ± 100 | <.001* | 446 ± 70 | 494 ± 60 | <.001<sup>⊗</sup> |
| S | 4 | 457 ± 62 | 209 ± 109 | <.001* | 464 ± 67 | 533 ± 89 | <.001<sup>⊗</sup> |
| S | 5<sup>†</sup> | 444 ± 46 | 465 ± 30 | .008<sup>⊗</sup> | 426 ± 73 | 416 ± 47 | .336 |
| S | 6<sup>†</sup> | 435 ± 40 | 426 ± 43 | .276 | 469 ± 60 | 458 ± 56 | .280 |
| S | 7 | 440 ± 62 | 221 ± 107 | <.001* | 504 ± 89 | 700 ± 173 | <.001<sup>⊗</sup> |
| S | 8 | 471 ± 63 | 185 ± 97 | <.001* | 488 ± 77 | 571 ± 98 | <.001<sup>⊗</sup> |
| S | 9<sup>†</sup> | 465 ± 48 | 465 ± 62 | .982 | 427 ± 74 | 452 ± 67 | .078 |
| S | 10 | 465 ± 64 | 188 ± 54 | <.001* | 439 ± 75 | 568 ± 108 | <.001<sup>⊗</sup> |

<sup>†</sup> Saline session\* Injection PSV < Control PSV (all $P < .050$ )⊗ Injection PSV > Control PSV (all $P < .050$ )

**Details of the peak saccade parameters during the MGS task for the SC inactivation experiments.** Each row contains an individual experimental session, and each column contains the details of that session. The column descriptions are as follows: “Monkey”: indicates which monkey the experiment was performed with; “Exp”: session number by animal (saline injection session are indicated with a dagger superscript); “Contralateral PSV Ctrl”: peak saccade velocity (in °/s) for correct trials in which the monkey made a saccade towards one of the three targets that are *contralateral* to the injected SC hemisphere during the *control* block; “Contralateral PSV Inject”: peak saccade velocity (in °/s) for correct trials in which the monkey made a saccade towards one of the three targets that are *contralateral* to the injected SC hemisphere during the *post-injection* block; “*P*-value”: two-tailed p-value from a non-parametric permutation test (5000 permutations) to assess whether mean PSV was lower post- vs pre-injection for the *contralateral* saccade trials; “Ipsilateral PSV Ctrl”: peak saccade velocity (in °/s) for correct trials in which the monkey made a saccade towards one of the three targets that are *ipsilateral* to the injected SC hemisphere during the *control* block; “Ipsilateral PSV Inject”: peak saccade velocity (in °/s) for correct trials in which the monkey made a saccade towards one of the three targets that are *ipsilateral* to the injected SC hemisphere during the *post-injection* block; “*P*-value”: two-tailed p-value from a non-parametric permutation test (5000 permutations) to assess whether mean PSV was lower post- vs pre-injection for the *ipsilateral* saccade trials.

**Extended Table 3:** DMC performance during SC inactivation experiments

| Monkey | Exp | Control perf.<br>($\mu \pm \sigma$) | Inact. perf.<br>($\mu \pm \sigma$) | Fisher-exact<br>odds ratio | Fisher-exact<br>*P*-value |
| --- | --- | --- | --- | --- | --- |
| N | 1<sup>†</sup> | 86.81 ± 3.91 | 85.27 ± 4.06 | 1.136 | .633 |
| N | 2 | 81.82 ± 4.31 | 53.15 ± 6.56 | 3.966 | <.001* |
| N | 3 | 83.64 ± 4.37 | 50.99 ± 5.21 | 4.912 | <.001* |
| N | 4 | 89.64 ± 3.77 | 56.08 ± 4.31 | 6.778 | <.001* |
| N | 5 | 80.06 ± 4.30 | 67.76 ± 4.29 | 1.910 | <.001* |
| N | 6 | 88.10 ± 3.70 | 73.25 ± 4.42 | 2.703 | <.001* |
| N | 7 | 89.97 ± 3.11 | 45.41 ± 6.61 | 10.785 | <.001* |
| N | 8<sup>†</sup> | 92.67 ± 3.09 | 89.43 ± 3.14 | 1.495 | .170 |
| S | 1<sup>†</sup> | 78.95 ± 4.73 | 82.44 ± 3.61 | 0.799 | .283 |
| S | 2 | 88.41 ± 3.61 | 64.03 ± 5.64 | 4.286 | <.001* |
| S | 3 | 87.84 ± 3.19 | 70.19 ± 8.79 | 3.068 | <.001* |
| S | 4 | 92.63 ± 3.48 | 56.52 ± 8.27 | 9.663 | <.001* |
| S | 5<sup>†</sup> | 86.05 ± 4.23 | 84.56 ± 4.19 | 1.126 | .716 |
| S | 6<sup>†</sup> | 93.95 ± 2.79 | 93.05 ± 2.13 | 1.159 | .662 |
| S | 7 | 95.74 ± 2.47 | 75.00 ± 6.63 | 7.485 | <.001* |
| S | 8 | 98.58 ± 1.38 | 75.82 ± 6.22 | 22.159 | <.001* |
| S | 9<sup>†</sup> | 92.46 ± 2.96 | 94.87 ± 2.45 | 0.663 | .248 |
| S | 10 | 97.24 ± 1.89 | 73.33 ± 5.59 | 12.818 | <.001* |

† Saline session
\* Injection perf. < Control perf. (all $P < .050$ )

**Details of performance on the DMC task for the SC inactivation experiments.** Each row contains an individual experimental session, and each column contains the details of that session. The column descriptions are as follows: “Monkey”: indicates which monkey the experiment was performed with; “Exp”: session number by animal (saline injection session are indicated with a dagger superscript); “Control perf.”: Mean ± SD accuracy (in %) for DMC trials during the control block; “Inact. perf.”: Mean ± SD accuracy (in %) for DMC trials during the post-injection block; “Fisher-exact odds ratio”: odds ratio for Fisher-exact test to compare performance on control vs. post-injection blocks.; “Fisher-exact *P*-value”: p-value for Fisher-exact test to compare performance on control vs. post-injection blocks.

## Notes

### Competing Interest Statement

The authors have declared no competing interest.

